# The PSD-95 interactome of hippocampal synapses potentiated in response to an in vivo learning task

**DOI:** 10.64898/2026.09.09.749100

**Authors:** Ajesh Jacob, Mariachiara Di Caprio, Ivan Arisi, Francesco Gobbo, Francesca Chiara Latini, Andrea Faraone, Lorena Zentilin, Ivonne Heinze, Silvia Marinelli, Alessandro Ori, Marco Mainardi, Antonino Cattaneo

**Affiliations:** Laboratory of Biology, Scuola Normale Superiore (Bio@SNS), Pisa, Italy; Present address: Arterra Bioscience SpA, Naples, Italy; European Brain Research Institute “Rita Levi-Montalcini” (EBRI), Rome Italy; Institute of Translational Pharmacology, National Research Council (IFT-CNR), Rome, Italy; Present address: Institute for Neuroscience and Cardiovascular Research and UK Dementia Research Institute, The University of Edinburgh, Edinburgh, United Kingdom; Molecular Medicine, International Centre for Genetic Engineering and Biotechnology, Trieste, Italy; Leibniz Institute on Aging - Fritz Lipmann Institute (FLI), Jena, Germany; Genentech Inc, South San Francisco, United States; Institute of Neuroscience, National Research Council (IN-CNR), Pisa, Italy; Department of Biomedical Sciences, University of Padua, Padua, Italy; Padua Neuroscience Center, Padua, Italy

**Author notes:** Equal contribution to this work (Mariachiara Di Caprio, Ivan Arisi). Antonino Cattaneo sadly passed away on June 28, 2026.

## Abstract

Changes in the molecular composition of synapses underlie the functional changes in synaptic strength after learning. These phenomena have been studied either via imaging of individual candidate proteins, or via the global analysis of the synaptic proteome, lumping together all synapses regardless of their activation or potentiation status. As a step towards the characterization of the proteome of synapses that have undergone potentiation following a learning paradigm, we have designed SynActive-PSD95-FLAG, a genetically encoded probe consisting of FLAG-tagged PSD- 95 endowed with activity-dependent local translation and expression enrichment at potentiating dendritic spines. SynActive-PSD95-FLAG was expressed via Adeno Associated Viral vectors (AAVs) in the hippocampus of mice which were exposed to contextual fear conditioning. Then, PSD95-FLAG immunoprecipitation from hippocampal extracts, followed by mass spectrometry, was used to characterize the changes in the PSD-95 interactome caused by in vivo synaptic potentiation. This revealed extensive rearrangements in the stoichiometry between PSD-95 and its direct and indirect interactors, highlighting proteins involved in peptide translation, including ribosomal components, and vesicle trafficking. Our results, obtained with a selectively expressed proteomic handle, provide new insight into the molecular substrates of hippocampal learning and a novel tool to study the proteome of potentiated synapses.

## Introduction

Dendritic spines have been under the spotlight of research on synaptic plasticity thanks to continuous progress in imaging methods and in the creation of probes for fluorescent protein tagging and Calcium imaging^1–4^. An increased volume of dendritic spine head represents the structural substrate for potentiation of synaptic transmission^5,6^ to the extent it is considered a proxy for synaptic potentiation^7,8^. A larger head corresponds to a wider postsynaptic membrane surface, required to accommodate more neurotransmitter receptors, as well as to a larger volume for soluble and scaffolding proteins. In this regard, relocalization of RhoA, Cdc42, CaMKII and glutamate receptor subunits has been shown to occur at individual spines following induction of structural potentiation^2,9,10^. These studies have captured the dynamics of individual molecules in great detail but cannot give a complete picture of the rearrangements in the synapse proteome that support synapse potentiation. In addition, these analyses have been mostly performed in cultured neurons or organotypic slices^2,3,5^, which complicates the extraction of molecular principles directly relevant for in vivo synaptic plasticity processes related to learning and memory.

In parallel to single-protein imaging studies, high-throughput proteomic methods for the global analysis of large populations of synaptic proteins from dissected brain samples have been set up. Initially, synaptosome preparations were combined with two-dimensional electrophoresis and mass spectrometry (MS) to identify a relatively small number of proteins^11^. Subsequently, isolation of the postsynaptic density fraction was combined with immunoprecipitation (IP) of the postsynaptic hub protein PSD-95^12^ and MS to qualitatively identify its interactome^13^, i.e. the set of direct and indirectly bound proteins. This approach was further refined with the creation of a knock-in transgenic mouse line expressing a double-tagged version of PSD-95 that was exploited for tandem affinity purification (TAP)^14^.

These studies have identified hundreds of components of the PSD-95 interactome and represent important steps towards the complete description of the entire postsynaptic proteome. On the other hand, these techniques are independent of the activation status of synapses and cannot be easily adapted to appreciate the modifications in the synaptome triggered by learning, which in physiological conditions take place in a relatively small subpopulation of synapses of a target brain area. To increase the anatomical resolution of synapse proteomics, laser capture microdissection has been successfully used to perform deep proteomic profiling of synapses from different hippocampal subregions^15^, but this approach cannot per se be exploited to sort synapses according to their potentiation state.

Towards this aim, we elaborated SynActive (SA), a genetically encoded tool designed to express reporters or actuators selectively at potentiated synapses^16–18^. SA exploits the properties of the 5’ and 3’ untranslated regions (UTRs) of the mRNA for the immediate-early gene *Arc*, namely its activity-dependent transport to dendrites close to the base of dendritic spines in a translationally repressed form, followed by activity-dependent translation in correspondence to dendritic spines undergoing synaptic potentiation^19,20^. We previously demonstrated that SA is able to restrict the expression of different fluorescent reporters and optogenetic probes specifically at potentiated synapses in vivo^16^.

In the present work, we exploited SA regulatory sequences to obtain in vivo state-dependent expression of a FLAG-tagged version of PSD-95 at potentiated synapses (SA-PSD95-FLAG).

The sequence encoding SA-PSD95-FLAG was packed into Adeno-Associated Viral vectors (AAVs) for delivery to and expression in the hippocampal CA1 area of mice exposed to contextual fear conditioning, followed by anti-FLAG IP and label-free tandem MS. In parallel, samples from mice constitutively expressing PSD95-FLAG were similarly processed. Comparative analysis of the two protein datasets allowed us to identify a large network of proteins whose direct or indirect interaction with PSD-95 increases in response to an associative learning behavioral task.

## Methods

### 1. Generation of constructs

All the plasmids for this study (**Suppl. Fig. 1**) were constructed by PCR amplifying each element and inserting it into the AAV vector pAAV-hSyn-EGFP-WPRE-pA (Addgene 50465). The 3×FLAG tag was synthesized as a gBlocks Gene fragment (Integrated DNA Technologies). The mouse PSD-95 coding sequence was amplified from PSD-95-TS:YSOG1 (Addgene 42226). The SARE-ArcMin promoter was synthesized as gBlocks Gene fragment, then the SARE enhancer fragment (-6,793 to -6,690; +1 denotes the transcription initiation site) was multiplexed to construct the E-SARE (5×SARE-ArcMin) synthetic promoter described in ^21^. The full-length Arc 5’ and 3’ untranslated regions (UTRs) was amplified from the SA-Ch plasmid described in ^16^. The polyadenylation sequence, consisting of the upstream sequence element and the full late polyadenylation signal sequence of SV40, was derived from pAAV-CW3SL-EGFP (Addgene 61463) and replaced the WPRE and bGHpA in the pAAV vector. All plasmids used were fully sequenced.

### 2. Preparation of adeno associated viral vectors (AAVs)

The custom-made recombinant AAVs used in this study were prepared by the AAV Vector Unit at the International Centre for Genetic Engineering and Biotechnology (ICGEB, Trieste, Italy; http://www.icgeb.org/avu-core-facility.html), as described previously^22^ with minor modifications. Briefly, infectious recombinant AAV vector particles were generated in HEK293T cells cultured in roller bottles by a cross-packaging approach whereby the vector genome was packaged into AAV capsid serotype 5 (pDP5, PlasmidFactory GmbH & Co. KG, Germany). Viral stocks were obtained by PEG precipitation and CsCl_2_ gradient centrifugation^23^. The physical titer of recombinant AAVs was determined by quantifying vector genomes (vg) packaged into viral particles by real-time PCR against a standard curve of a plasmid containing the vector genome^24^, and corresponded to 5.4×10^11^ vg/ml for Cnst-PSD95-FLAG and to 4.7×10^12^ vg/ml for SA-PSD95-FLAG. The AAV2/PHP.eB CAG::tdTomato AAV was purchased from Addgene (56462; 1.0×10^13^ vg/ml).

### 3. Primary cultures

#### a. Primary neuronal cell culture

Primary hippocampal neurons were extracted from postnatal day (P) 0 B6;129 mice as follows: after surgical isolation of the hippocampus, tissue was triturated in cold Ca^2+^-free Hank’s balanced salt solution (HBSS, Invitrogen) with 100 U/mL penicillin, 0.1 mg/mL streptomycin, followed by trypsin digestion (0.1%, 20 min, 37°C) and inactivation in Dulbecco’s modified Eagle’s medium (DMEM; Invitrogen) supplemented with 10% fetal bovine serum (FBS, Euroclone) and 100 U/mL DNase. After centrifugation (5 min, 1000 rpm), the cell pellet was resuspended in Neurobasal-A medium (Invitrogen) supplemented with 4.5 g/L D-glucose, 10% FBS, 2% B27 (Invitrogen), 1% L-glutamine, 1mM sodium pyruvate and 12.5 mM glutamate. Neurons were plated on poly-D-lysine-coated glass coverslips in 24-well plates at a density of 1.0×10^5^ neurons per well (total volume for each well was 500 μl) and placed in a humidified incubator (37°C, 5% CO_2_).

On the first day in vitro (DIV1), the medium was replaced with Neurobasal-A medium supplemented with 2% B27, 1% L-glutamine, 10 mg/ml gentamicin and 12.5 mM glutamate. On DIV2, 2.5 mM cytosine β-D-arabinofuranoside (AraC, C1768, Merck) was added to the medium to suppress proliferation of non-neuronal cells. From DIV3 onwards, glutamate was removed from the medium described above. The medium was refreshed every other day by replacing half of the total volume.

#### b. In vitro infection

On DIV7-8, cells were infected with either AAV2/5 SA-PSD95-FLAG or Cnst- PSD95-FLAG at a multiplicity of infection (MOI) of 1.0×10^4^ vg/cell.

#### c. Pharmacological treatments

Experiments were performed at DIV14. Neurons underwent glycine-strychnine chemical LTP (cLTP) according to the following protocol: (i) the entire medium for each well was collected and kept at 37°C in a water bath; (ii) 500 μL of Extracellular Solution (ECS), containing, in mM, NaCl 140, glucose 33, HEPES 25, KCl 5, CaCl_2_ 1.3, and, in μM, picrotoxin 50, strychnine 1, tetrodotoxin 0.5, pH 7.4, was added to each well, and cells were incubated for 30 min at 37°C; (iii) half of the ECS was removed and kept at 37°C, cLTP was induced with 250 μL fresh ECS + 200 μM glycine for 3 min; (iv) after cLTP induction, the ECS was replaced with 250 μL fresh ECS + 250 μL conditioned ECS solution from (iii) for 30 min. Finally, the ECS was removed and replaced with conditioned medium from (i). Three hours after cLTP induction, cells were washed with PBS and fixed with paraformaldehyde (PFA; 4%, 10 min, RT).

### 4. In vitro immunofluorescence and imaging

Cells were fixed with PFA (2% w/v in PBS) and permeabilized with PBS containing 0.1% Triton X- 100 and 2.5% bovine serum albumin (BSA) for 7 min. Then, samples were blocked with PBS containing 5% BSA for 1 h, followed by a 2.5 h incubation in PBS containing 2.5% BSA, in addition to primary antibodies against FLAG (made in rabbit, 1:1,600 dilution, Cell Signaling Tech. 14793) and MAP-2 (made in mouse, 1:2,000 dilution, Abcam ab5392) at RT, rocking. After washing in PBS containing 2.5% BSA, the secondary antibodies, anti-rabbit conjugated to AlexaFluor 488 (1:400, Invitrogen A32731) and anti-mouse conjugated to AlexaFluor 555 (1:400, Invitrogen A28180), were dissolved in the same solution and added for 1 h at RT, rocking. After a final wash in PBS, coverslips were mounted on glass slides using Fluoroshield (Sigma-Aldrich).

Images were analyzed using Fiji/ImageJ^25^. FLAG-immunoreactive puncta decorating MAP-2-labeled neurites were manually counted. The length of analyzed neurites was measured to obtain the linear density of FLAG-immunoreactive puncta/mm.

### 5. Stereotaxic injections

Stereotaxic microinjections were performed by adapting the procedures described in ^26,27^. C57BL/6J male mice (2 months old) were used for all experiments. Mice were anesthetized by intraperitoneally injecting zoletil-xylazine (80-10 mg/kg). Body temperature was kept constant throughout the procedures at 36°C with a closed-loop temperature control system (Harvard Apparatus). Before surgery, the scalp was treated with topical anesthetic (2.5% lidocaine ointment). Two holes were opened in the skull using a microdrill to insert heat-pulled micropipettes fabricated from calibrated glass capillaries (Merck BR708707) at stereotaxic coordinates −2.0 mm AP, ±1.9 mm ML relative to bregma, and −1.4 mm DV, corresponding to the dorsal CA1 area of the hippocampus. An oil microinjector (CellTram 4r Oil, Eppendorf) was employed to deliver 1 μl of AAV solution at a rate of 0.1 μl/min. The concentration of AAVs was adjusted to deliver equal amounts of each preparation (5.4×10^8^ vg). Only for immunofluorescence experiments, either SA-PSD95-FLAG or Cnst-PSD95- FLAG AAVs were mixed 9:1 with CAG::Tomato AAV. After the injection procedure, the micropipette was held in place for 10 min to prevent spillover of the solution during subsequent retraction. Finally, the wound was cleaned and sutured with 5/0 vicryl non-absorbable thread (Ethicon). Mice received oral paracetamol (0.1 mg/g) for 3 days; 21 days were allowed for complete recovery and to achieve steady-state expression of the AAV-borne gene constructs.

### 6. Contextual fear conditioning

All behavioral experiments were performed using a fear conditioning chamber controlled via the AnyMaze software (Ugo Basile). A testing chamber with electrified grid floor, was placed into a white sound-attenuating box (48.5×38.5×48.5 cm) equipped with loudspeaker, a ventilation fan and a camera mounted on the ceiling. A high-contrast pattern was attached to the walls of the chamber to characterize the context (conditioning stimulus, CS) where the electric shock (unconditioned stimulus, US) was to be delivered. Every day, mice were acclimated to the testing room for 30 min. On day 1 (habituation phase), mice injected with the SA-PSD95-FLAG AAV were exposed to the conditioning context for 5 min. On day 2 (learning phase) to the room, mice were exposed to the same conditioning context, where they were subjected to 5 foot shocks (0.5 mA, 1 s, 25 s intershock interval) during the first 3 minutes; then, 2 minutes were allowed for further exploration of the context.

### 7. Histological fixation

Three hours after contextual fear conditioning, mice were overdosed with i.p. chloral hydrate (10% w/v), then the heart was exposed and the left ventricle was cannulated using a 27G needle connected to silicone tubing passing through a peristaltic pump (Gilson). After cutting the wall of the right atrium, transcardial perfusion started at a flow rate of 18.0 ml/min. First, 25 ml of ice-cold PBS, pH 7.4, were passed to clear tissues from blood, followed by at least 35 ml of PFA (4% w/v in 0.1 M phosphate buffer, PB, pH 7.4). Brains were extracted from the skull and postfixed in 4% PFA for 2- 6 hours. Samples were finally transferred to 30% w/v sucrose in 0.1 M PB and maintained at 4°C for at least 3 days, or until they sank to the bottom of the tube, for cryoprotection^28^.

### 8. Immunofluorescence and imaging of brain sections

Coronal sections (50-μm thickness) comprising the hippocampus were prepared using a sliding microtome (Leica SM2010R), then treated with blocking solution (10% NGS, 0.3% Triton X-100 in PBS) for 2 h, followed by incubation with primary antibodies (1:1,600 rabbit FLAG F7425, Merck Life Science; 1:300 guinea pig Homer-1b/c, Synaptic Systems 160025), diluted in PBS with 1% NGS, 0.1% Triton, overnight at 4°C, rocking. After washing once in PBS with 0.3% Triton X-100, and twice in PBS, 10 min, room temperature (RT), rocking, sections were incubated with secondary antibody solution (1:400 goat anti-rabbit Alexa Fluor-488 Thermo Fisher A-11008, 1:400 goat anti-guinea pig Alexa Fluor-647, Thermo Fisher A-21450) diluted in PBS with 1% NGS and 0.1% Triton X-100 for 2 h, RT, rocking. Then, sections were washed again thrice, as above, then mounted on microscope slides using Vectashield H-1000 medium. Images were acquired using a Zeiss LSM 900 confocal microscope (63×, N.A. 1.4, Airyscan mode). The number of FLAG- and Homer-1b/c-immunoreactive puncta along CA1 stratum radiatum dendrites was manually counted, with the investigator blind to experimental groups, and dendrite length measured to calculate the density of puncta/mm.

### 9. Tissue dissection, protein extraction, immunoprecipitation, and Western blotting

Three hours after contextual fear conditioning, mice were subjected to cervical dislocation, and brains were quickly extracted from the skull. After a quick wash in ice-cold PBS, brains were placed on a Petri dish filled with ice, then hippocampi were dissected out and immediately stored at -80°C. For each sample, hippocampi from 2 or 3 mice injected with SA-PSD95-FLAG or Cnst-PSD95-FLAG, respectively, were pooled to obtain enough material for the subsequent immunoprecipitation and homogenized in 500 μl/animal of ice-cold DOC buffer, containing 50 mM Tris pH 9.0, 50 mM NaF, 1mM Na_3_VO_4_, 20 μM ZnCl, 1% sodium deoxycholate, 1 tablet/10 ml Complete Mini Protease Inhibitor Cocktail (Roche 04693116001) and 1 tablet/10 ml PhosSTOP phosphatase inhibitor (Roche 11699695001). After a 1-h incubation on ice, the extracts were clarified by centrifugation at 13,000g for 30 min at 4°C. These “input” samples were incubated with anti-FLAG M2 magnetic beads (Sigma M8823) (1 mg/ml; 50 μl/animal) for 2 h at 4°C with constant agitation. Beads were washed three times in DOC buffer without deoxycholate, and immunoprecipitated proteins were eluted from the beads by heating in 50 μl/animal of elution buffer, containing 100 mM Tris-HCl pH 8.0 and 4% SDS for 10 min at 90°C. The total protein concentration in each sample was quantified using Micro BCA Protein Assay Kit (Thermo Fisher Scientific 23235). Aliquots from each sample, containing 2-10 μg of total proteins, were subjected to SDS-PAGE on 10% acrylamide gels, transferred to Protran 0.45 nitrocellulose membranes (GE Healthcare Life Science 10600114). After blocking in 5% skimmed milk (Bio-Rad 1706404) for 1 h, blots were incubated with primary antibodies overnight at 4°C with gentle shaking. Blots were then washed thrice in TBST (Tris-buffered saline, 0.1% Tween-20) and incubated with horseradish peroxidase (HRP)-conjugated secondary antibodies for 1 h, RT, rocking. The primary and secondary antibodies used were: mouse monoclonal FLAG M2 (1:1,000, Sigma F1804), rabbit polyclonal GluN2A/B (1:1,000, Synaptic Systems 244003), rabbit monoclonal GluA1 (1:1,000, Cell signaling 13185), goat polyclonal CaMKII (1:500, Thermo Fisher Scientific PA5- 19128), rabbit monoclonal Arc (1:1,000, Cell Signaling 65650), rabbit monoclonal S6 (1:1,000, Cell Signaling 2217), rabbit polyclonal Histone H3 (1:1,000, Abcam 1791), mouse IgGκ-binding protein-HRP (1:2,000, Santa Cruz 516102), mouse anti-rabbit IgG-HRP (1:2,000, Santa Cruz 2357), mouse anti-goat IgG-HRP (1:2,000, Santa Cruz 2354). Following three washes in TBST, chemiluminescence signals from HRP were detected using SuperSignal West Femto Maximum Sensitivity Substrate (Thermo Fisher Scientific 34093) using a ChemiDoc MP Imaging system (Biorad). Following image acquisition, the optical density of the bands of interest was quantified by adapting the approach described in ^29^. The following normalization operations were performed for each sample: (i) [(PSD95-FLAG IP blot) / (PSD95-FLAG input blot)], (ii) (protein of interest IP blot / PSD95-FLAG IP blot). Normalized values were further normalized to the mean of Cnst-PSD95-FLAG for each blot to allow pooling of different experimental replicas.

### 10. Mass spectrometry

#### a. Sample preparation

Input and immunoprecipitation samples were supplemented with the same volume of 2× SDS lysis buffer (100 mM HEPES and 2% w/v SDS). Samples were sonicated using a Bioruptor Plus (Diagenode) for 15 min (10 cycles: 1 min on, 30 sec off, 20 °C) using the high setting, then boiled for 10 min at 95°C, followed by a second round of sonication (as before). Samples were spun down at 18,407g for 1 min and the lysate supernatant transferred to fresh tubes. Cysteine residues were reduced by adding 200 mM dithiothreitol (DTT) to a final concentration of 10 mM (incubated for 15 min at 45°C). Samples were then alkylated by adding 200 mM iodoacetamide to a final concentration of 15 mM (incubated for 30 min at RT in the dark). Proteins were then precipitated with 4 volumes ice-cold acetone to 1 volume sample and left overnight at -20°C. Samples were then centrifuged at 20,817g for 30 minutes, 4°C. After removal of the supernatant, the precipitates were washed twice with 300 µL 80% v/v acetone (ice-cold). After each wash step, the samples were vortexed, then centrifuged again for 10 minutes at 4°C first, then 5 minutes. The pellets were then allowed to air-dry before being dissolved in digestion buffer (3 M urea in 0.1 M HEPES, pH 8, 1:100 w/w of LysC, Wako 125-05061) and incubated for 4 h at 37°C, shaking at 650 rpm. Then, samples were diluted 1:1 with HPLC water (bringing urea to 1.5 M) and were incubated with 1:100 w/w trypsin (Promega V5280) for 16 h at 37°C, shaking at 650 rpm. Digests were then acidified with 10% trifluoroacetic acid (TFA) and then desalted with Waters Oasis HLB µElution Plate 30 µm in the presence of a slow vacuum. In this process, columns were conditioned with 3×100 µL Oasis solvent B (80% v/v acetonitrile; 0.05% v/v formic acid, FA) and equilibrated with 3×100 µL Oasis solvent A (0.05% v/v FA in milliQ water). Samples were loaded, washed 3 times with 100 µL solvent A, and then eluted into PCR tubes with two lots of 50 µL each of solvent B. Eluates were dried down with the speed vacuum centrifuge at 45°C in vacuum-aqueous mode. Dried peptides were next dissolved in reconstitution MS buffer A (5% v/v acetonitrile, 0.1% v/v TFA in water) and analyzed by mass spectrometry.

#### b. Data acquisition

Peptides were separated in trap/elute mode UltiMate 3000 UPLC system (Thermo Fisher Scientific) fitted with a trapping (Waters nanoEase M/Z Symmetry C18, 5 μm, 180 μm × 20 mm) and an analytical column (Waters nanoEase M/Z Peptide C18, 1.7 μm, 75 μm × 250 mm). Solvent A was water, 0.1% FA and solvent B was 80% (v/v) acetonitrile, 0.08% FA. One µl of the sample (∼1 μg) was loaded with a constant flow of solvent A at 5 μl/min onto the trapping column. Trapping time was 6 min. Peptides were eluted via the analytical column with a constant flow of 0.3 μl/min. During the elution, the percentage of solvent B increased in a nonlinear fashion from 0–48% in 96 min. Total run time was 115 min. The outlet of the analytical column was coupled directly to a Q exactive HF (Thermo Fisher Scientific) using the Proxeon nanospray source. The peptides were introduced into the mass spectrometer via a Pico-Tip Emitter outer diameter 360 μm × inner diameter 20 μm, tip 10 μm (New Objective) heated to 300 °C, and a spray voltage of 2.2 kV was applied. The capillary temperature was set at 300°C. The radio frequency ion funnel was set to 50%. For data-independent acquisition (DIA), full scan mass spectrometry (MS) spectra with mass range 350–1650 m/z were acquired in profile mode in the Orbitrap with resolution of 120,000 FWHM. The default charge state was set to 3+. The filling time was set at maximum of 60 ms with limitation of 3×10^6^ ions. DIA scans were acquired with 34 mass window segments of differing widths across the MS1 mass range. Higher collisional dissociation fragmentation (stepped normalized collision energy; 25.5, 27, and 30%) was applied and MS/MS spectra were acquired with a resolution of 30,000 FWHM with a fixed first mass of 200 m/z after accumulation of 3×10^6^ ions or after filling time of 40 ms (whichever occurred first). Data were acquired in profile mode. For data acquisition and processing of the raw data Xcalibur 4.1 (Thermo Fisher Scientific) and Tune version 2.9 were used.

#### c. Raw data analysis

Acquired data were processed using Spectronaut Professional v15.10 (Biognosys AG). For library creation, the DIA raw files were searched with Pulsar (Biognosys AG) against the mouse UniProt database (*Mus musculus*, entry only, release 2016_01) with a list of common contaminants appended, using default settings. For library generation, default BGS factory settings were used. DIA data were searched against this spectral library using BGS factory settings, except: Proteotypicity Filter = Only Protein Group Specific; Major Group Quantity = Median peptide quantity; Major Group Top N = OFF; Minor Group Quantity = Median precursor quantity; Minor Group Top N = OFF; Data Filtering = Qvalue sparse; Normalization Strategy = Local normalization; Row Selection = Automatic.

### 11. Bioinformatic analysis

Each entry of the SA and Cnst IP datasets was normalized to the total content of the PSD-95 bait in the corresponding experimental replica, by calculating normalized expression (NE) = [log_2_(iBAQ entry) - log_2_(iBAQ PSD-95)]; each value was averaged among the experimental replicas. Then, differential protein expression was measured as SA - Cnst = (mean NE for SA) - (mean NE for Cnst).

Following the approach of ^34^, the statistical significance of SA vs. Cnst differences was assessed using t-test with Benjamini-Hochberg correction for multiple comparisons. Heatmaps with dendrograms were created by the ggplot2 R package^35^. Pathway analysis of protein lists was computed by the clusterProfiler R package^36^, using the KEGG, Reactome and Gene Ontology Biological Processes databases of functional gene categories, using as background for enrichment all genes annotated to each database in the dataset Org.Mm.eg.db for *Mus musculus*. The ClusterProfiler tool was employed to perform Fisher’s exact test, and FDR multiple testing correction was used to test pathway overrepresentation. Network analyses were performed using the STRING database^37^ through the Cytoscape^38^ application, using a high confidence criterion (0.7) and restricting retrieved nodes to those physically interacting with PSD-95.

## Results

### 1. Design and in vitro validation of gene constructs for state-dependent PSD-95 interactomics

To characterize the PSD-95 interactome of potentiated synapses, we designed a construct for the expression of a FLAG-tagged version of PSD-95 as a proteomic bait enriched at potentiated synapses. To target the expression of the PSD-95-FLAG mRNA to dendrites of activated neurons and enrich it at or close to potentiated synapses, we employed the SynActive regulatory sequences^16^, thus obtaining SA-PSD95-FLAG. Activity-dependent transcription of SA-PSD95-FLAG was achieved under the control of the synthetic, activity-dependent E-SARE promoter^21^, followed by the transport of mRNA into the active dendrites enabled by the Arc-derived SA sequences. The activity-dependent translation of the dendritic PSD95-FLAG mRNA close to potentiated synapses would then allow enrichment of the PSD95-FLAG protein into the potentiating spine. As our overall goal was to define the changes in the PSD-95 interactome taking place at potentiated synapses in comparison to baseline synaptic activity, we reasoned that constitutive expression of the same PSD95-FLAG probe via the human synapsin promoter would be an ideal term of comparison. We therefore prepared two constructs, named E-SARE::SynActive-PSD95-FLAG (SA-PSD95-FLAG) and synapsin::PSD95-FLAG (Cnst-PSD95-FLAG), to express PSD95-FLAG at potentiated synapses or in a constitutive and potentiation-independent fashion, respectively (**Fig. 1A**).

**Figure 1.**
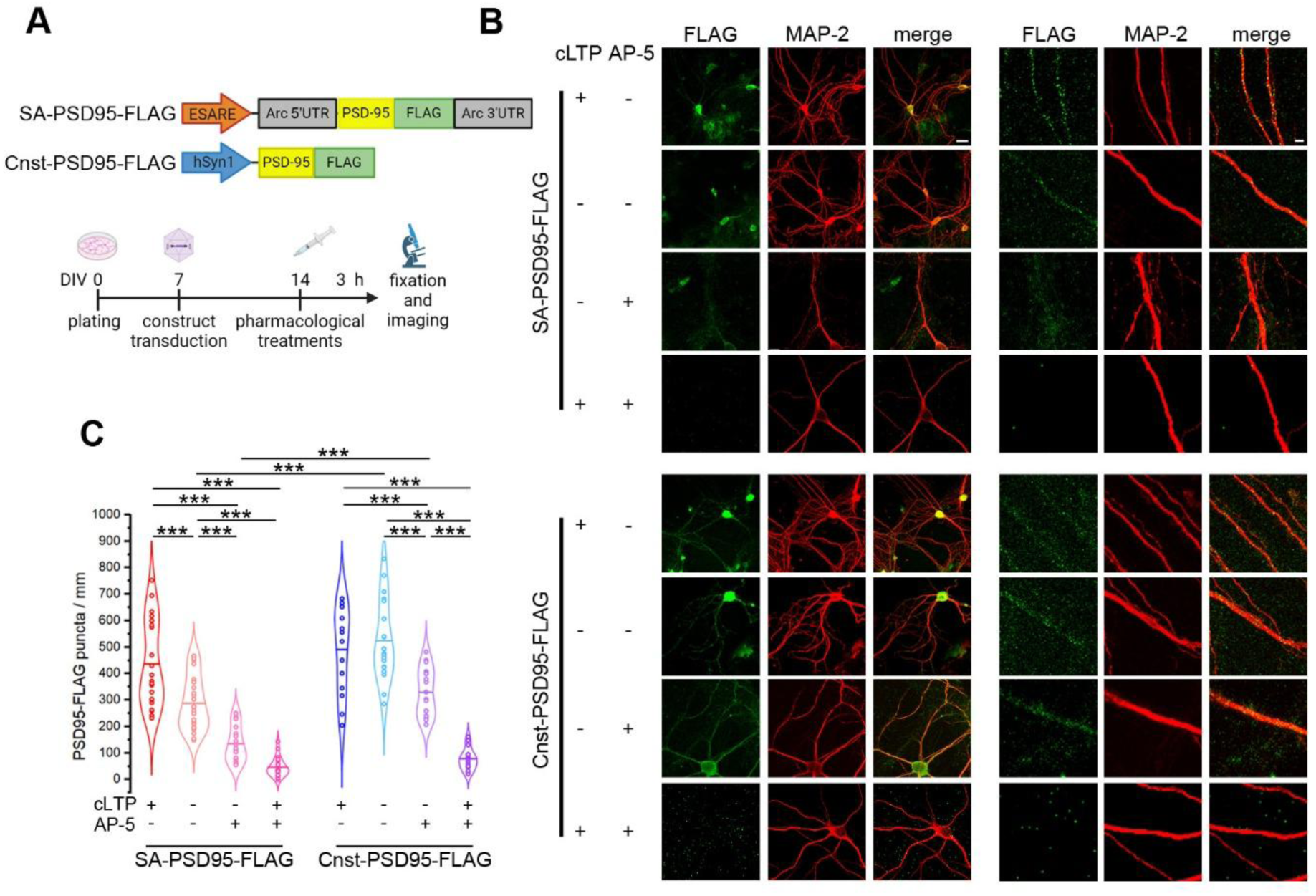
SynActive allows the expression of PSD95-FLAG in a potentiation-dependent pattern in primary neuronal cultures. **A)** Scheme of construct design and timeline of experiments. **B)** Representative high- and low-magnification images showing the expression of PSD95-FLAG expressed in SynActive (SA) or constitutive (Cnst) modes in MAP-2-positive neurites. Scale bars are 40 μm and 5 μm for left and right panels, respectively, **C)** Quantification of the density of FLAG- positive puncta reveals a significant increase in neurons expressing SA-PSD95-FLAG after induction of chemical LTP (cLTP) while neurons expressing Cnst-PSD95-FLAG (i.e., constitutive expression of PSD95-FLAG) fail to show such modulation by cLTP. Moreover, neurons expressing SA-PSD95- FLAG show lower baseline levels of FLAG-positive puncta and increased susceptibility to NMDA receptor blockade (ANOVA-2 construct × treatment F_3,160_=8.410, p<0.001, followed by Bonferroni post-hoc test, ***p<0.001; SA-PSD95-FLAG+cLTP, n=22, SA-PSD95-FLAG-cLTP, n=29, SA- PSD95-FLAG-cLTP+AP-5, n=20, SA-PSD95-FLAG+cLTP+AP-5, n=21, Cnst-PSD95-FLAG+cLTP, n=14, Cnst-PSD95-FLAG-cLTP, n=19, Cnst-PSD95-FLAG-cLTP+AP-5, n=19, Cnst-PSD95- FLAG+cLTP+AP-5, n=17 dendrites).

The expression patterns of SA-PSD95-FLAG and Cnst-PSD95-FLAG were first analyzed in primary hippocampal neuronal cultures. Neurons expressing SA-PSD95-FLAG exposed to chemical long- term potentiation (cLTP) displayed an increased density of FLAG-immunoreactive puncta along MAP-2-positive dendrites, in comparison to vehicle-treated neurons expressing the same construct. On the other hand, neurons infected with Cnst-PSD95-FLAG showed no significant difference in the number of FLAG-immunoreactive puncta between cLTP and vehicle. In basal conditions (i.e. treatment with vehicle), the density of FLAG-immunoreactive puncta in neurons expressing Cnst- PSD95-FLAG was higher than for neurons expressing SA-PSD95-FLAG, indicating, as expected, that the latter construct is expressed only in a subset of the whole synaptic population. Upon cLTP induction, the number of FLAG-positive puncta in neurons expressing SA-PSD95-FLAG increased greatly, and no significant difference compared with neurons expressing Cnst-PSD95-FLAG was detected (**Fig. 1B-C**). This could be explained by saturation of synaptic potentiation in this in vitro system. For both groups, treatment with the NMDA receptor (NMDAR) antagonist AP-5, either alone or in combination with cLTP induction, resulted in a significant reduction of PSD95-FLAG-positive puncta. In neurons expressing SA-PSD95-FLAG, AP-5 treatment alone caused a 53.3% reduction in the density of FLAG-positive puncta in comparison to the basal level, while in Cnst-PSD95-FLAG- expressing neurons the reduction by AP-5 treatment alone was 37.2%. This suggests a higher sensitivity of the SA-PSD95-FLAG puncta signal to NMDAR activity than the Cnst-PSD95-FLAG signal, further supporting dependency of SA-construct expression upon synaptic potentiation (**Fig. 1B-C**). Appropriate controls confirmed the immunoreactivity specificity **(Suppl. Fig. 2**).

These data provide in vitro evidence that the SynActive-based approach can be successfully used to enrich the expression of a proteomic probe at synapses undergoing activity-dependent potentiation.

### 2. In vivo expression of the SA-PSD95-FLAG proteomic probe in the mouse hippocampus following fear conditioning

The results from in vitro experiments prompted us to investigate the in vivo expression of PSD95- FLAG in synapses potentiated by an in vivo learning paradigm using contextual fear conditioning (CxFC), a hippocampus-dependent paradigm widely used to study associative learning and memory^39,40^.

SA-PSD95-FLAG or Cnst-PSD95-FLAG-encoding constructs were delivered to the CA1 area of the mouse hippocampus using Adeno-Associated Viral vectors (AAVs). Three weeks later, learning-dependent synaptic potentiation of hippocampal synapses was triggered by exposing mice to the association phase of CxFC. Three hours after CxFC, mice underwent histological fixation or fresh tissue dissection (see the workflow of **Fig. 2A**).

**Figure 2.**
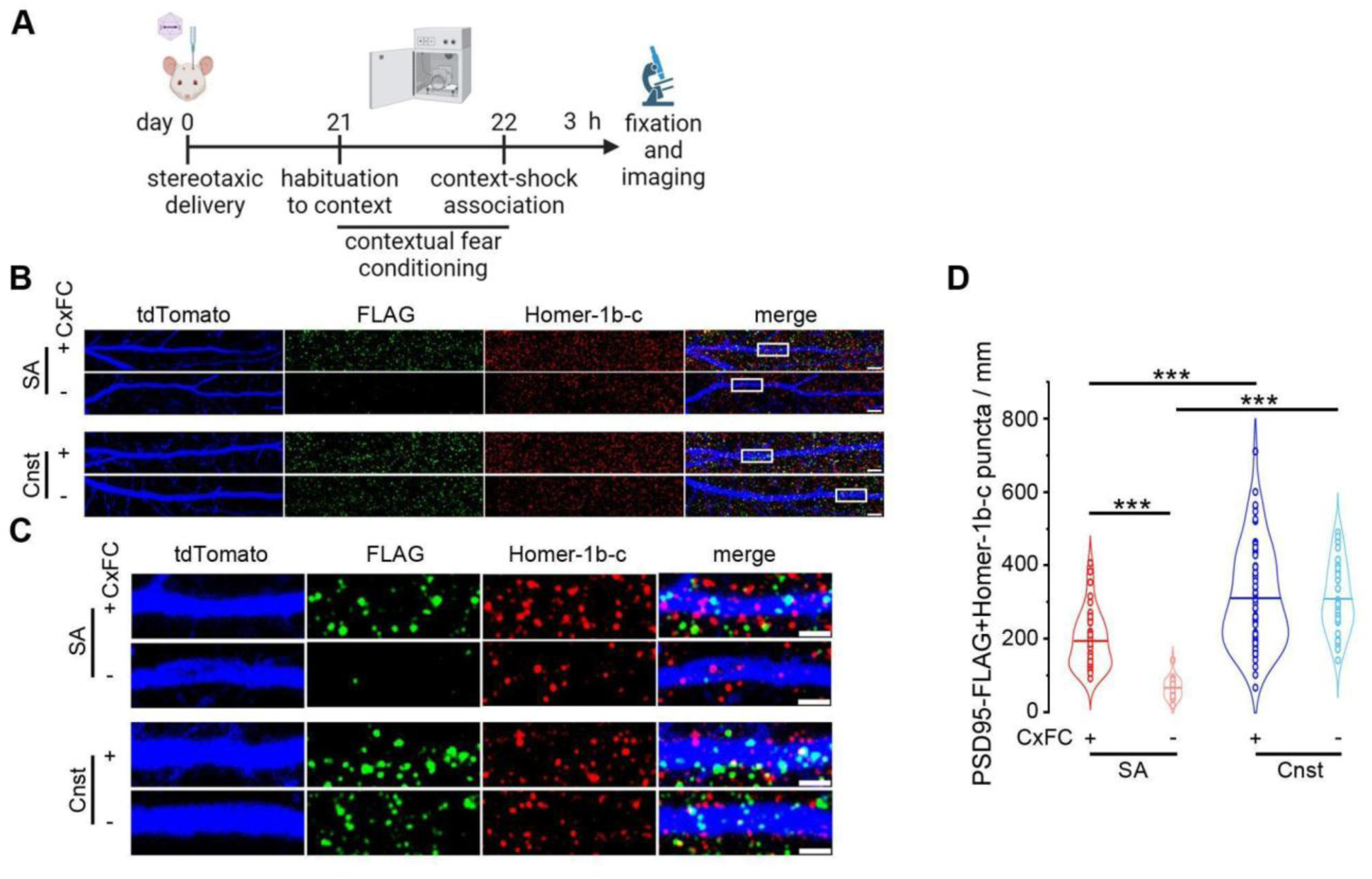
SynActive-driven expression of PSD95-FLAG in potentiated synapses in CA1 hippocampal neurons in vivo after Contextual Fear Conditioning. **A)** Timeline for in vivo experiments. **B)** Representative images showing the colocalization of PSD95-FLAG puncta with Homer-1b-c puncta on dendrites of CA1 neurons, labeled via sparse co-infection with a synapsin-tdTomato AAV, in SA-PSD95-FLAG (SA)- or Cnst-PSD95-FLAG (Cnst) AAV-injected mice. Scale bars are 5 μm. **C)** Magnified views of dendrite sections framed in white rectangles on panel B. Scale bars are 2 μm. **D)** Quantification of the density of FLAG-positive puncta reveals a significant increase in neurons expressing SA-PSD95-FLAG (SynActive) after exposure of mice to contextual fear conditioning (CxFC +) in comparison to mice kept in their home cages (CxFC -). On the other hand, the density of FLAG-positive puncta is unaffected by learning in neurons from mice expressing Cnst- PSD95-FLAG (constitutive) (ANOVA-2 construct × behavioral task F_1,158_=11.268 p<0.001, followed by Bonferroni post-hoc test, ***p<0.001; SA+CxFC, n=44; SA-CxFC, n=18; Cnst+CxFC, n=66; Cnst- CxFC, n=33 dendrites).

First, we checked the expression pattern of PSD95-FLAG by immunofluorescence. To facilitate the detection of PSD95-FLAG-positive puncta in dendrites, an AAV expressing the tdTomato red fluorescent protein under the control of the synapsin promoter (synapsin::tdTomato) was co-injected with either SA-PSD95-FLAG or Cnst-PSD95-FLAG AAVs, with titer adjusted to obtain sparse labeling of dendritic arbors **(see Methods; Fig. 2B**).

It is noteworthy that the in vivo expression of SA-PSD95-FLAG was totally restricted to dendrites, and no labeling was detected in the cell bodies, thus showing an even higher localization precision in comparison to primary neuronal cultures **(Suppl. Fig. 3).** This matches what we previously observed^16^, and is likely related to the fact that the regulation of the reporter expression by the Arc regulatory sequences is more preserved in the intact in vivo circuitry than in cultured neurons.

Quantification of the density of SA-PSD95-FLAG-immunoreactive puncta colocalizing with the postsynaptic marker Homer-1b-c revealed that exposure to CxFC of SA-PSD95-FLAG AAV-injected mice doubled the density of synaptically localized FLAG-immunoreactive puncta in comparison to mice kept in their homecage (**Fig. 2C-D**). On the other hand, no modulation of FLAG-immunoreactive puncta was observed in Cnst-PSD95-FLAG AAV-injected mice after exposure to CxFC (Fig. 2D). Of note, the density of FLAG-immunoreactive puncta in mice expressing SA-PSD95-FLAG was markedly lower than in mice expressing Cnst-PSD95-FLAG, both after CxFc and in homecage conditions (**Fig. 2D**). The total density of Homer-1b-c puncta was not significantly different among the four experimental groups, indicating that expression of SA-PSD95-FLAG or Cnst-PSD95-FLAG per se did not perturb synapse density **(Suppl. Fig. 4)**. Control experiments confirmed the specificity of FLAG and Homer-1b-c immunoreactivity **(Suppl. Fig. 5)**.

### 3. Using the SA-PSD95-FLAG proteomic probe to characterize the changes in the PSD-95 interactome following in vivo synaptic potentiation

Having established the enriched expression of the Synactive-controlled SA-PSD95-FLAG proteomic bait at potentiated synapses in vitro and in vivo, we sought to use this tool to characterize state-dependent PSD-95 interactomics in hippocampal neurons in vivo, following exposure to the learning phase of CxFC.

To obtain a global picture of the changes in the PSD-95 interactome following learning-induced synaptic potentiation, protein extracts from the hippocampi of SA- and Cnst-PSD95-FLAG- expressing mice were subjected to co-immunoprecipitation (IP) using anti-FLAG beads and analyzed by label-free tandem MS. We chose to compare samples from mice expressing SA-PSD95- FLAG exposed to the learning phase of CxFC with samples from mice expressing Cnst-PSD95- FLAG kept in their home cages, which would represent conditions of behavior-induced synaptic potentiation and of baseline synaptic activity in the absence of novel stimuli, respectively. Mice that did not receive any AAV injection (CTRL) were also included in our analysis (see below).

Quantification of the total content of PSD-95 (namely, endogenous PSD-95 plus PSD95-FLAG) in total protein extracts (i.e., input to IP) showed that constitutive expression of PSD95-FLAG driven by the synapsin promoter resulted in a small, albeit significant increase of total PSD-95 in comparison to CTRL mice. On the other hand, SynActive-controlled expression did not result in a significant difference in total PSD-95, compared to CTRL mice, nor to Cnst mice **(Suppl. Fig. 6A)**. This shows that the contribution of AAV-encoded PSD95-FLAG to the total pool of endogenous PSD-95 is marginal and agrees with unaltered synapse density (**Suppl. Fig. 4**). Moreover, expression of neither construct had behaviorally detectable consequences, as we observed no significant difference in the freezing time between Ctrl, Cnst and SA groups **(Suppl. Fig. 6B)**. Comparison of the PSD-95 levels in FLAG IPs yielded a radically different picture, with both Cnst and SA groups showing a significantly higher content than the background, represented by the Ctrl group. Moreover, the PSD-95 content in IP samples from the Cnst group was higher than in the SA group **(Suppl. Fig. 6A**), in keeping with the higher density of synaptic puncta tagged under constitutive expression conditions compared to potentiation-dependent labeling (**Fig. 2D**).

We then analyzed the global mass spectrometry data from SA- or Cnst-PSD95-FLAG IP samples from mice that had undergone CxFC.

The correlation index dendrogram Input samples were grouped together, regardless of the experimental group they belonged to. On the contrary, IP samples belonging to SA and Cnst groups were segregated from their respective Input material, as well as from Ctrl IPs which, instead, correlated with Ctrl Inputs. Consistently, IP samples from SA and Cnst groups were segregated from those obtained from Ctrl mice **(Suppl. Fig. 7**).

This initial global assessment of samples was followed by analytical elaboration of the MS dataset, which contained a total of 3,892 identified proteins **(Suppl. Table 1)**. We undertook a four-step sequential filtering procedure of this dataset to exclude proteins showing: i) non-specific binding to IP beads, ii) consistently low frequency of detection across samples iii) no significant enrichment in IP samples from mice expressing either SA-PSD95-FLAG or Cnst-PSD95-FLAG compared to IP samples from mice not expressing FLAGged PSD95 (i.e., Ctrl), (iv) annotation lacking a known synaptic or dendritic localization **(Suppl. Fig. 8**).

In the first filtering step, we looked for proteins which might have been included in the dataset because of non-specific binding to anti-FLAG conjugated agarose beads, which would manifest as a significant enrichment in IP versus Input samples obtained from CTRL mice, i.e. not expressing the FLAGged IP bait. This comparison yielded 244 proteins, which were classified as "non-specific binders" and were therefore excluded from subsequent analyses **(Suppl. Table 2)**. The second filtering step consisted in verifying that a given protein was represented at least 3 times (out of a total of 5 experimental replicas) in both SA and Cnst groups; 162 proteins failed to meet this criterion and were, thus, excluded from the subsequent comparative analysis **(Suppl. Table 3)**.

This left us with a total of 3,489 proteins, which underwent a third filtering step to exclude 69 proteins that did not show a statistically significant change in their abundance in IP samples from SA or Cnst groups compared to IP samples from the Ctrl group (**Suppl. Table 4**).

The final filtering step excluded from further analysis proteins which were annotated for non-synaptic localization and not for synaptic, nor dendritic localization, nor synapse-associated localization (**Suppl. Table 5**; **Fig. 3A**), resulting in a final dataset of 2,609 proteins (**Suppl. Table 6**).

**Figure 3.**
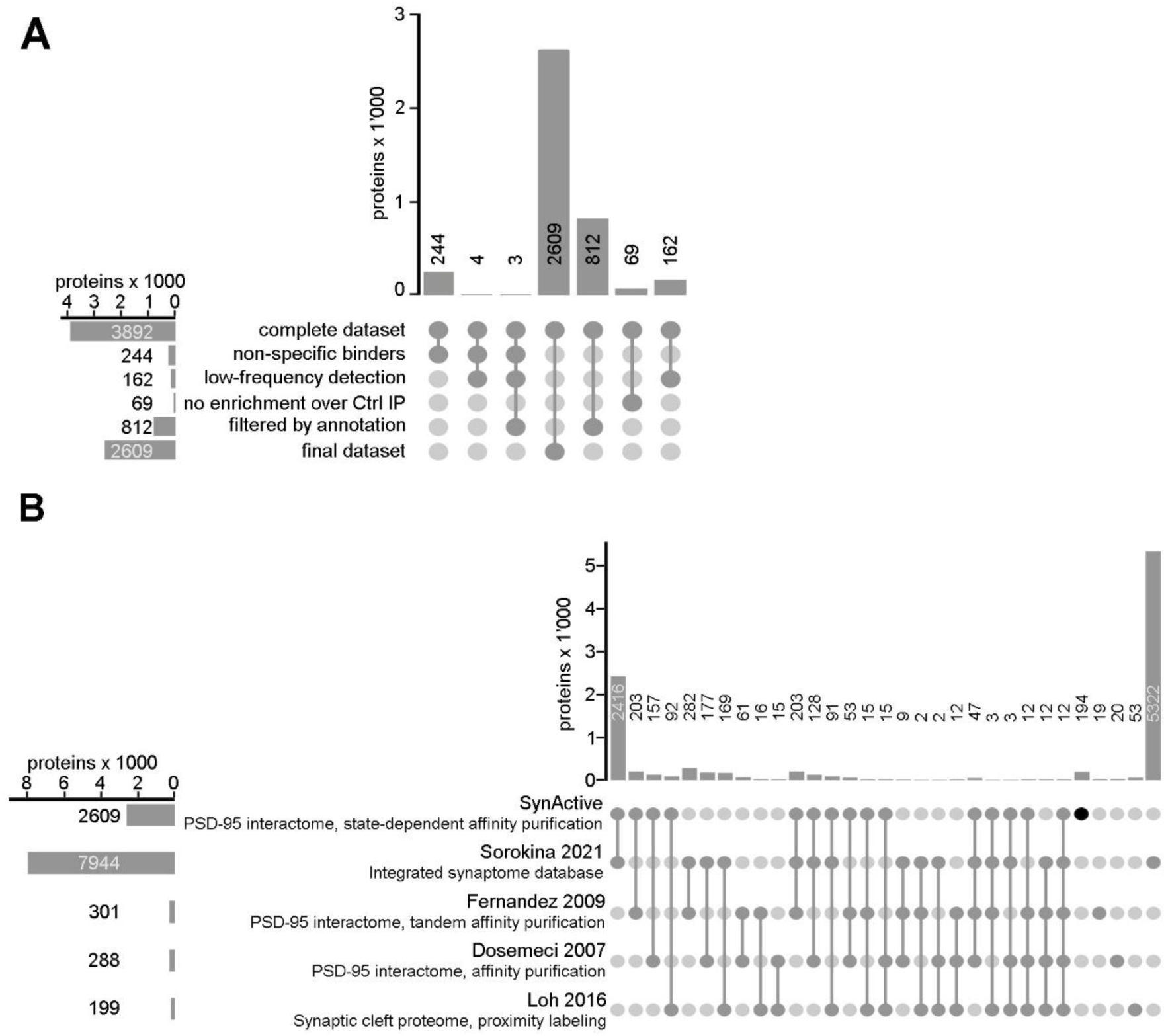
Global analysis of the PSD-95 interactome from potentiated and basally active synapses. **A)** Upset plot diagram showing filtering operations on the initial list of 3,892 PSD-95 interactors identified by mass spectrometry. Sewage of entries showing significant enrichment also in IP samples from mice not expressing PSD95-FLAG (non-specific binders), unreliable detection by MS in either experimental group (low detection), no significant enrichment compared to Ctrl IP, and no annotation for synaptic localization (filtering by annotation) yielded a dataset of 2,609 proteins. **B)** Intersections of our 2,609-protein set with previously published PSD-95 interactomes^13,14^, synaptic cleft proteome^41^ and a comprehensive synaptome database^42^ highlights a high level of overlap with published data, along with 194 newly identified proteins. Of note, 12 core proteins are contained in all datasets, including AMPA and NMDA receptors subunits, neurexin-1, and neuroligin 3.

To frame our dataset in the context of previously published synaptic proteome analyses, we performed a comparison with 4 main published datasets^13,14,41,42^ (**Fig. 3B**). This highlighted 194 new entries which are present only in our dataset **(Suppl. Table 7)**. The remaining entries showed a high overlap (93%, 2,416 out of 2,609 proteins) with the comprehensive overall synaptome database compiled by Sorokina et al. ^42^ **(Suppl. Table 8)**. Of note, the concordance with the PSD-95 interactomes published in Fernandez et al.^14^ and Dosemeci et al.^13^ was about 67% and 55% (203 out of 301 proteins and 157 out of 288 proteins, respectively; **Suppl. Table 8**). Finally, 46% of the proteins (92 out of 199) contained in the excitatory synaptic cleft proteome of Loh et al.^41^ were also contained in our dataset **(Suppl. Table 8)**. This comparison also highlighted the existence of a core, formed by 12 proteins, which is common to the 4 published datasets and to ours (**Fig. 3B; Suppl. Table 9)**, namely Cacng2/stargazin, AMPA and NMDA receptor subunits, kainate receptor isoform 2, Lrrtm1, neuroligin-3, and neurexin-1.

We performed a similar comparison using the “non-specific binders” list **(Suppl. Fig. 9**). Only 15 “non-specific” entries were exclusively found in our experimental dataset, while 229 entries were also part of the Sorokina et al.^42^ database **(Suppl. Table 10)**. On the other hand, 13, 15, and 6 entries were contained in the intersections between the non-specific binders list and the Fernandez et al.^14^, Dosemeci et al.^13^ and Loh et al.^41^ lists, respectively **(Suppl. Table 10)**.

### 4. Comparative analysis of the PSD-95 interactome from potentiated versus basally active synapses

The SA-PSD95-FLAG and Cnst-PSD95-FLAG IP datasets overlapped in terms of qualitative protein composition and were strongly correlated (**Fig. 4A**). Indeed, synaptic potentiation is likely to mainly involve changes in the quantity of a given protein localized at synapses, in line with recent deep proteomic profiling of different hippocampal synaptic types showing that their functional diversity arises from changes in the abundance of shared components^15^. To gain insight into this point, we normalized the abundance of each protein to the total content of the PSD95-FLAG bait in the corresponding samples, then ranked them according to the difference in their abundance between SA and Cnst groups (**Fig. 4B**). Finally, we statistically compared the two datasets following the approach of Rayaprolu et al.^34^, and found that 2,581 entries showed a significantly increased abundance in the PSD-95 interactome in response to learning-induced synaptic potentiation **(Suppl. Table 11)**. These included all the 12 core proteins common to ours and published datasets **(Suppl. Table 9; Fig. 4C**). Moreover, among the proteins showing the highest difference between SA and Cnst groups, we found known PSD-95 interactors, including AMPA receptor subunits, PSD-95- associated scaffolds, activity-regulated kinases and signal transducers, Rho GTPases, in addition to fractalkine/Cx3cl1 and the C3 complement element (**Fig. 4D**). Of note, Lrrtm1 and Lrrtm2, which were previously used to identify the synaptic cleft proteome^41^, were also enriched in the potentiated synapse (SA) IP group (**Fig. 4C and Suppl. Table 11**).

**Figure 4.**
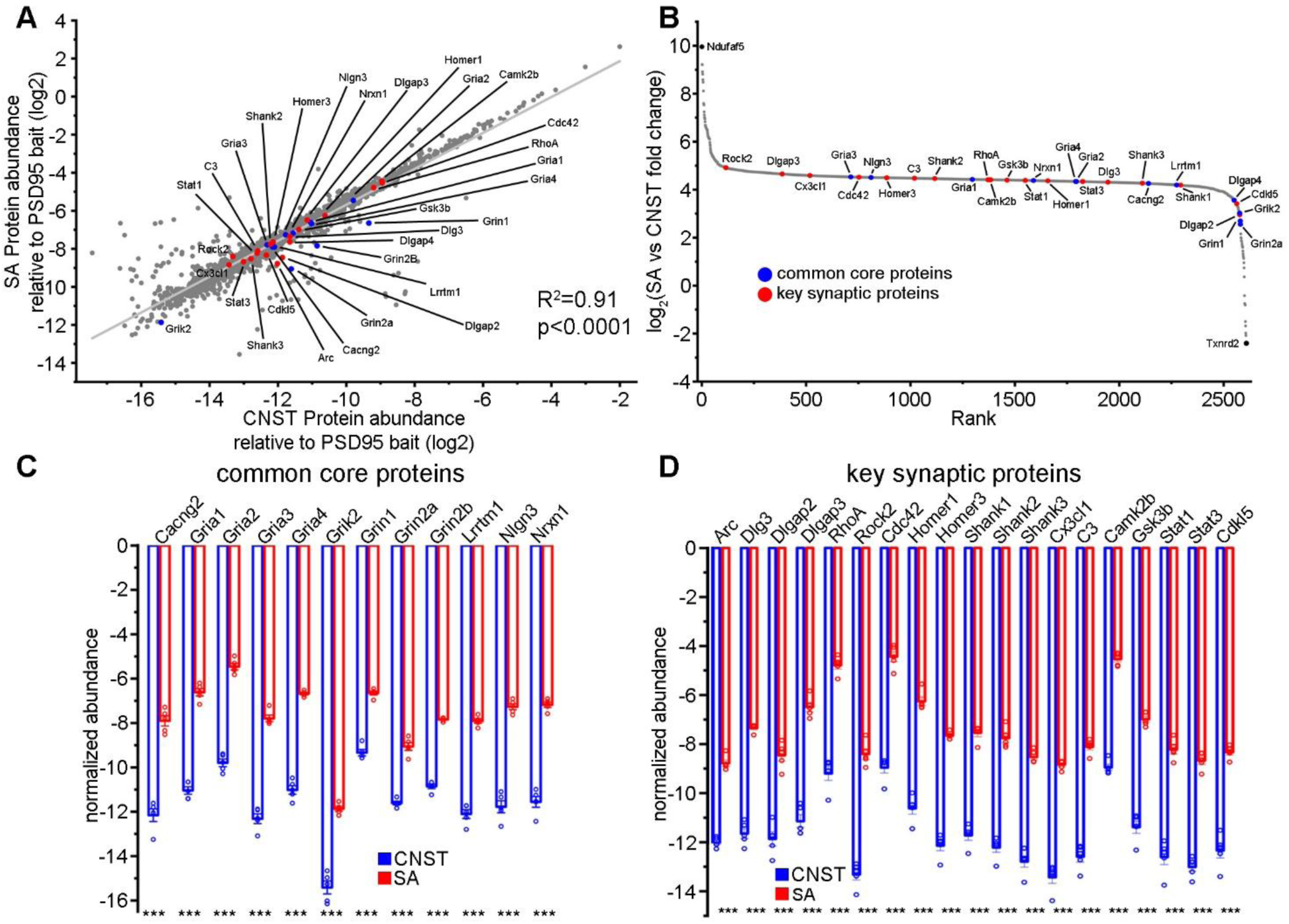
Understanding the PSD-95 interactome of potentiated synapses. **A)** Scatter plot showing statistically significant positive correlation between the SA and CNST datasets. **B)** Rank plot showing differentially expressed proteins between SA and CNST groups. **C)** Histograms showing increased interaction with PSD-95 of the 12 core proteins common to our and published datasets (n=5 for SA and Cnst, ***p<0.001). **D)** Histograms showing increased interaction between key synaptic proteins and PSD-95 upon synaptic potentiation (n=5 for SA and Cnst, ***p<0.001). Negative values for protein abundance are due to the log_2_(protein abundance)-log_2_(PSD-95 abundance) normalization.

Then, we validated the MS data via Western blotting analysis of a set of candidate PSD-95 interactors, as an independent method to quantify the expression of known PSD-95 interactors in the hippocampus of animals injected with SA-PSD95-FLAG vs Cnst-PSD95-FLAG. Both IP samples and the corresponding input material were analyzed. The overall amount of PSD95-FLAG that could be retrieved from hippocampi expressing SA-PSD95-FLAG was lower in comparison to Cnst- PSD95-FLAG samples (**Fig. 5A**), in keeping with the property of SynActive-controlled proteins to show enriched expression selectively at the subset of synapses undergoing potentiation. This result is also consistent with the lower density of FLAG-immunoreactive puncta in hippocampal neurons observed in mice expressing SA-PSD95-FLAG compared to Cnst-PSD95-FLAG (**Fig. 2**). On the other hand, the yield of IP was comparable between hippocampi expressing SA-PSD95-FLAG or Cnst-PSD95-FLAG, as demonstrated by comparable IP/Input material ratios (**Fig. 5A**). Blots were then probed with antibodies specific for key interactors of PSD-95 involved in synaptic plasticity, namely NMDA receptor subunits 2A-B (GluN2A-B; SA vs Cnst MS fold change - FC - 2.55 for GluN2A and 3.01 for GluN2B, p<0.0001), AMPA receptor subunit 1 (GluA1; FC 4.43, p<0.0001), calmodulin-dependent kinase IIα (CaMKIIα; FC 4.28, p<0.0001), and activity-regulated cytoskeletal protein (Arc; FC 3.25, p<0.0001)^14^ (**Fig. 5B**). The optical density ratio (IP/Input material) between the blot bands corresponding to each interactor and PSD-95 was lower in samples from mice expressing SA-PSD95-FLAG compared to Cnst-PSD95-FLAG for all proteins tested (**Fig. 5C**). This supports the capability of SA-controlled PSD95-FLAG to retrieve both direct and indirect PSD-95 interactors from the small subset of synapses undergoing potentiation, while constitutive expression of PSD95-FLAG leads to retrieving proteins from the bulk of hippocampal synapses, regardless of their potentiation state. However, when the amount of each immunoprecipitated protein was normalized to the corresponding PSD95-FLAG band, we detected a highly significant enrichment in IP samples from SA-PSD95-FLAG mice in comparison to Cnst-PSD95-FLAG (**Fig. 5D**). This finding suggests an increased stoichiometric ratio between PSD-95 and its interactors upon synaptic potentiation. We used histone 3 (H3) as a negative control for possible nuclear contaminants in our samples and found that H3 was virtually undetectable in IP samples, with no significant difference between SA-PSD95-FLAG and Cnst-PSD95-FLAG (**Fig. 5B,E**). Finally, we analyzed ribosomal protein S6 (rpS6; FC 4.43, p<0.0001), which can be involved in local protein translation at potentiating synapses^43,44^. Also in this case, we found a lower enrichment and a higher stoichiometric ratio with PSD-95 in IP samples from SA-PSD95-FLAG-expressing mice compared to Cnst-PSD95- FLAG (**Fig. 5F**).

**Figure 5.**
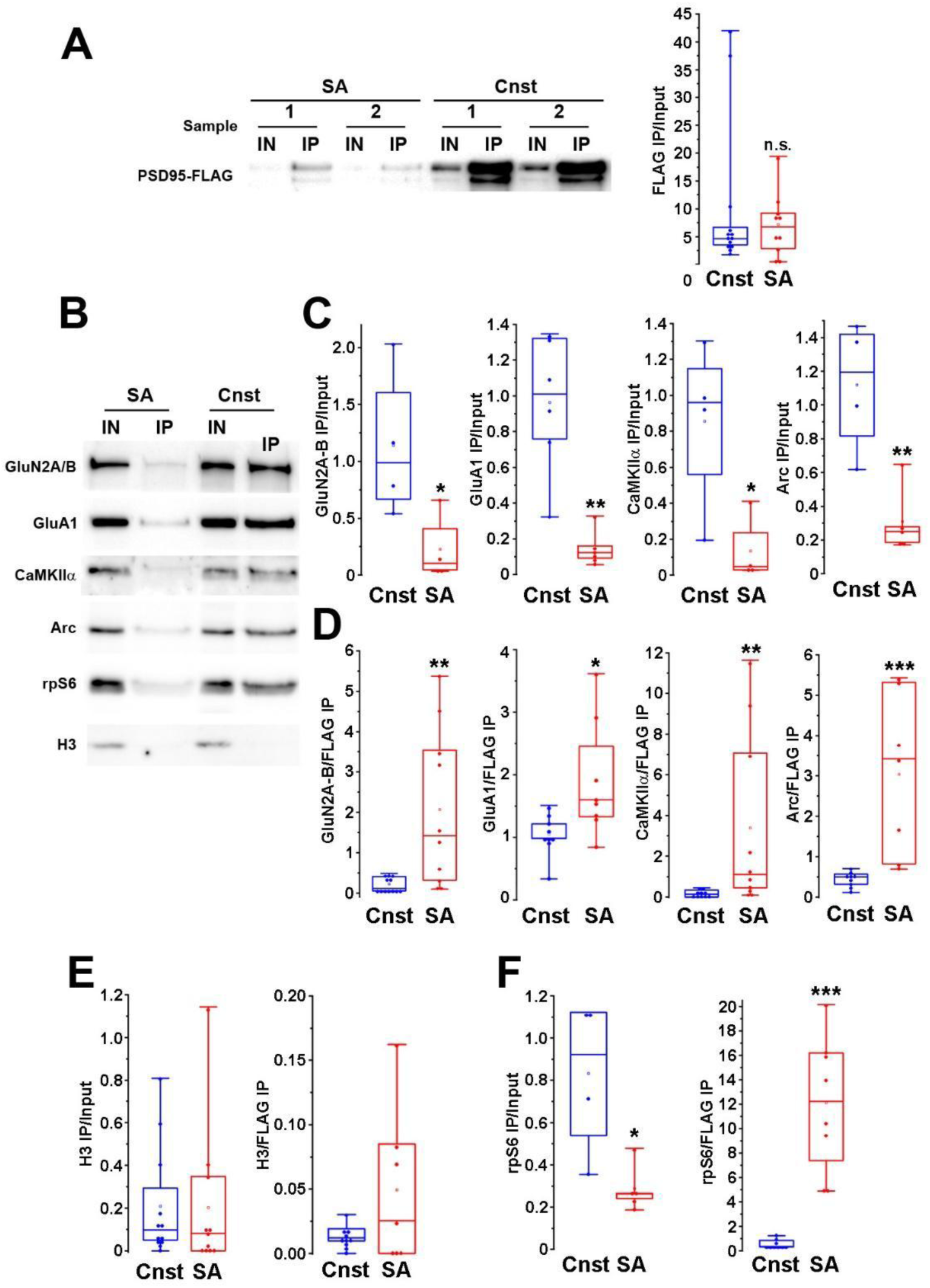
Immunoprecipitation of SynActive- and constitutive-PSD95-FLAG after fear conditioning recovers known interactors of PSD-95. **A)** Representative blots and quantification showing comparable enrichment for PSD95-FLAG in IP samples from mice expressing SA-PSD95- FLAG (SA) or Cnst-PSD95-FLAG (Cnst) with respect to Input material (Cnst, n=13; SA, n=11; Mann-Whitney U test, p=0.926). **B)** Representative blots containing IP and Input samples probed with antibodies for synaptic and non-synaptic proteins. **C)** Lower IP/Input material ratios for known PSD- 95 interactors in SA-PSD95-FLAG samples than in Cnst-PSD95-FLAG samples (GluN2A/B, Cnst, n=4; SA, n=4; *p=0.044; GluA1, Cnst, n=9; SA, n=8; **p=0.004; CaMKIIα, Cnst, n=4; SA, n=4; *p=0.029; Arc, Cnst, n=4; SA, n=5; **p=0.004; Mann-Whitney U test for all comparisons). **D)** Increased stoichiometry of interaction between PSD-95 and its known interactors in SA-PSD95- FLAG samples than in Cnst-PSD95-FLAG samples (GluN2A/B, Cnst, n=12; SA, n=10; **p=0.004; GluA1, Cnst, n=9; SA, n=8; *p=0.020; CaMKIIα, Cnst, n=10; SA, n=10; **p=0.004; Arc, Cnst, n=9; SA, n=7; ***p=0.001; Mann-Whitney U test for all comparisons). **E)** No significant enrichment in histone H3 (H3) in SA-PSD95-FLAG and Cnst-PSD95-FLAG IP samples (Cnst, n=12; SA, n=11; Manny-Whitney U test, p=0.404). **F)** Lower IP/Input ratio and higher stoichiometry of interaction with PSD-95 in SA-PSD95-FLAG samples compared to Cnst-PSD95-FLAG samples (IP/Input, Cnst, n=4; SA, n=5, Mann-Whitney U test, *p=0.016; IP/FLAG, Cnst, n=9; SA, n=8, Mann-Whitney U test, *p<0.001).

These experiments demonstrate that SA-PSD95-FLAG expression in vivo is sensitive to plasticity induced by learning and is synaptically localized. Moreover, IP using anti-FLAG magnetic beads of SA-PSD95-FLAG protein extracts can isolate the small amount of PSD-95 and its direct and indirect interactors contained in potentiated synapses.

### 5. Network analysis of the identified PSD-95 interactors enriched by synaptic potentiation

To gain insight into the relationships among the PSD-95 interactors affected by synaptic potentiation, we performed a network analysis on the 2,609-protein list obtained from the initial dataset filtering (**Suppl. Fig. 8**; **Suppl. Table 6**) by selecting those including in their GO Cellular Component annotation the terms "synapse", "synaptic", "dendrite", "dendritic" (**Suppl. Table 12**). The proteins fulfilling these criteria form a main network with 513 nodes, which were color-coded according to their function (**Fig. 6; Suppl. File 1)**. This network had a mean connectivity degree of 9.123, in stark contrast with the low connectivity (0.947 – 1.290) shown by random networks generated using the same 513 nodes **(Suppl. Fig. 10**). When we calculated the percentage of total nodes showing a given degree of connectivity (i.e., number of network edges), we found that most proteins had between 1 and 7 connections. Interestingly, proteins involved in membrane trafficking / synaptic vesicle cycle and synaptically or dendritically localized ribosomal components clusters formed two strikingly visible clusters with high connectivity, ranging from 15 to 25 and 39 to 45 edges, respectively **(Suppl. Fig. 11)**. The 44 ribosomal proteins we identified are a subset of those that can be retrieved by a standard query in UniProt **(Suppl. Fig. 12**; **Suppl. Table 13)**.

Fifty-four direct interactors of PSD-95 enriched at potentiated synapses corresponded to high-connectivity nodes of the network, mainly functioning as scaffolds (Shank1-2-3, Dlgap3-4), AMPA and NMDA receptor subunits (Gria1-4, Grin1, 2a-b), and signal transducers (Rho GTPases, CaMKII) **(Suppl. Fig. 13)**.

**Figure 6.**
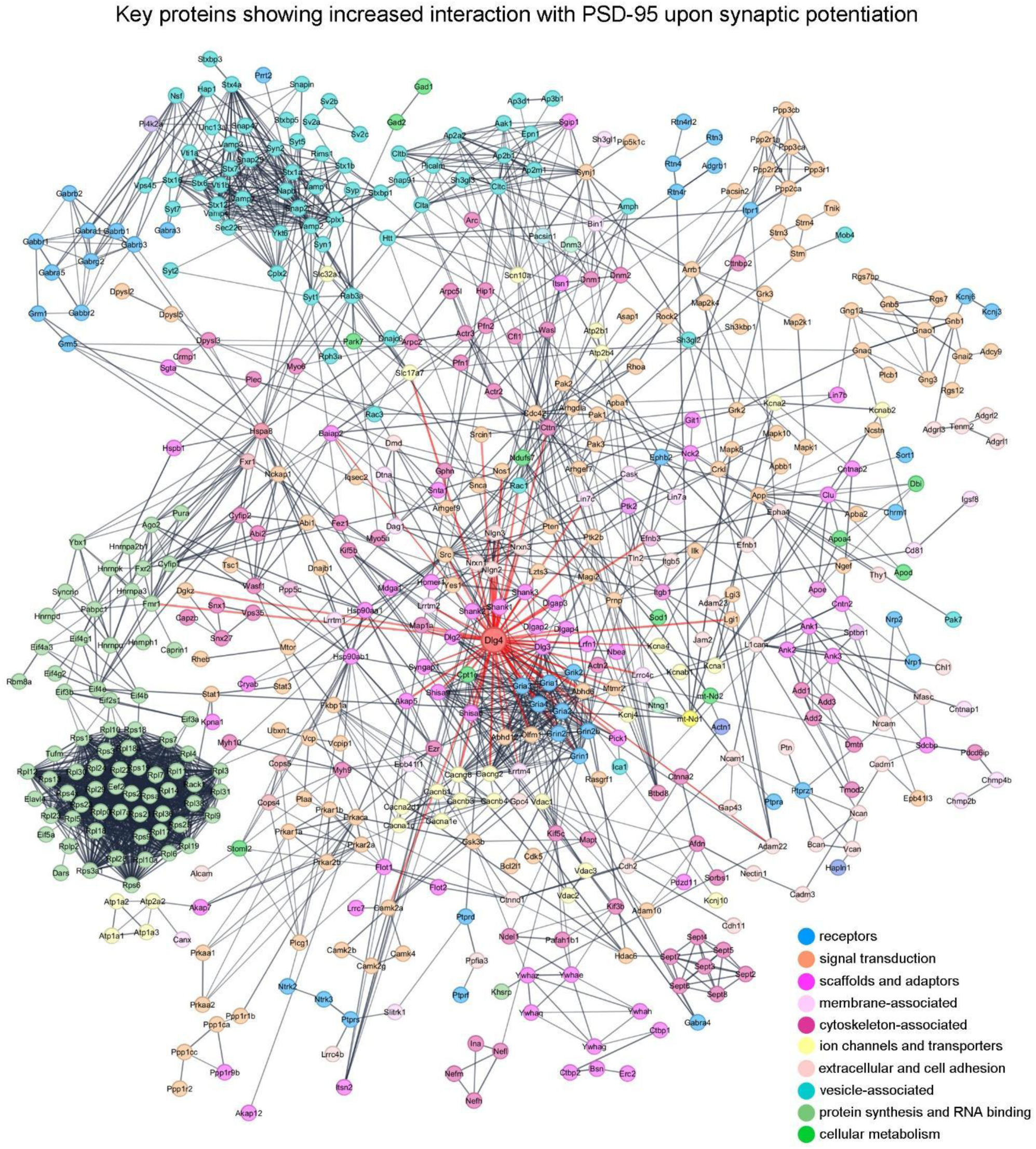
Network analysis of the PSD-95 interactors enriched at potentiated synapses. Network of proteins with significantly higher interaction with PSD-95 at potentiated synapses and previous annotation for synaptic or dendritic localization. Each node was colored according to its main cellular function. Edges connecting PSD-95 to its direct interactors are highlighted in red.

Pathway enrichment analysis provided an overall view of the protein families found to be enriched in the PSD-95 interactome at potentiated synapses. GO Biological Process entries showing significant false discovery rate (FDR) and high strength contained not only terms directly associated with synapse structure, but with vesicle-mediated transport, protein translation, and energy metabolism (**Fig. 7A**, **Suppl. Table 14)**. Consistently, significant GO Cellular Components and Molecular Function contained terms related to synapse structure and plasticity, protein translation, regulation of cytoskeleton, and mitochondrial activity **(Suppl. Fig. 14, Suppl. Tables 15 -16**). Relevant KEGG terms contained “Ig SF CAM signaling” (i.e., intercellular contact), "glutamatergic synapse", and "pathways of neurodegeneration" (**Fig. 7B**, **Suppl. Table 17)**. Finally, Reactome yielded high-strength, significant terms related to protein translation, receptor signaling and energy metabolism (**Fig. 7C**, **Suppl. Table 18)**.

**Figure 7.**
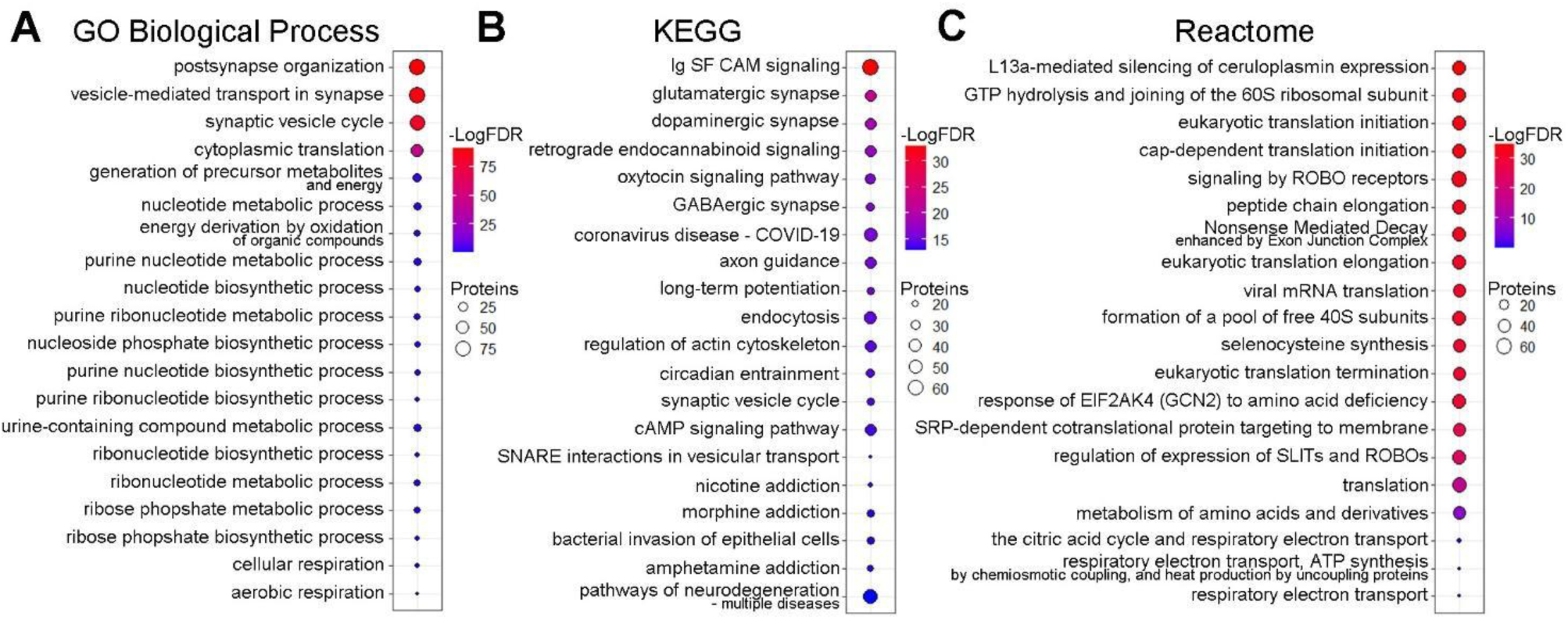
Gene ontology and pathways enriched in the PSD-95 interactome of potentiated synapses. **A, B, C)** Gene ontology (GO) Biological Process, KEGG and Reactome analyses for the nodes of the network shown in Fig. 6, ranked according to their odds ratio (see also **Suppl.** Fig. 14).

These results indicate that synaptic potentiation is supported not only by proteins primarily involved in synapse organization and synaptic transmission, but also by recruitment of additional elements supporting protein synthesis and trafficking.

## Discussion

The molecular composition of the excitatory postsynaptic dendritic spines changes dynamically as a function of their activity and potentiation state, due to concomitant activity-dependent processes: traffic of proteins into and out of the synaptic compartment^1^, local protein synthesis^45^, and degradation^46,47^.

Despite significant progress in the characterization of the synapse proteome^13,14,41,42^, no studies have directly tackled the problem of defining the molecular fingerprint associated with synaptic potentiation in vivo, due the lack of suitable experimental methods. All current methods to study the synaptic proteome face the problem of cellular and synaptic anatomical and functional heterogeneity. Ideally, one would need to engineer the expression of a proteomic bait selectively in a potentiated synapse, to pull down and identify the proteins from that synapse and not from other synapses of the same neuron that have not undergone potentiation. Towards this aim, we exploited SynActive (SA), a genetically encoded tool designed to express reporters or actuators selectively at (or close to) potentiated synapses^16–18^. We previously demonstrated that SA is able to restrict the expression of different fluorescent reporters and optogenetic probes specifically at potentiated synapses in vivo^16^. Here, we describe a method based on the SA strategy, that allows the expression of a proteomic^21^ bait (SA-PSD95-FLAG) specifically at potentiated synapses in vivo. The SA-PSD95-FLAG proteomic bait is a FLAG-tagged version of PSD-95 scaffold protein expressed under the transcriptional control of the E-SARE activity-dependent promoter^21^ and under the translational control of the Synactive 5’ and 3’ UTR regulatory sequences^16^.

Using this method, we report the identification of the PSD-95 interactome of hippocampal synapses after potentiation induced by fear conditioning.

The results represent a first proof of principle towards the systematic large-scale identification of proteins differentially expressed in the subset of all synapses undergoing learning-related long-term potentiation (**Suppl. Fig. 15**). or potentiation status (i.e., status unknown), including also synapses potentiated by endogenous synaptic plasticity processes (i.e., not related to specific behavioral stimuli or tasks).

We used Adeno Associated Viral vectors (AAVs) to express PSD95-FLAG in the CA1 area of mice, which were then exposed to contextual fear conditioning (CxFC), which is known to trigger functional^48^ and structural^49,50^ potentiation of CA1 synapses. Exposure to the learning phase of CxFC increased the number of FLAG-immunoreactive puncta on apical dendrites of CA1 neurons in mice expressing SA-PSD95-FLAG.

The constitutive expression of PSD95-FLAG via AAV-mediated transduction does not cause potential overexpression-related alterations^51–53^. Even though mass spectrometry (MS) could detect a small increase in the total amount of PSD-95 in protein extracts from animals constitutively expressing PSD95-FLAG, this exogenous protein was expressed in the expected punctate pattern and in close apposition with its interactor Homer-1 in both constitutive and SA conditions. Moreover, the behavioral response (i.e., freezing) to CxFC was comparable among the Cnst-PSD95-FLAG, SA-PSD95-FLAG and Ctrl (i.e., not expressing any recombinant gene construct) groups, supporting the conclusion that gene constructs did not cause overexpression-related alterations.

On the other hand, the large difference (4.33 log_2_ units) in the PSD-95 content observed in IP samples from Cnst and SA groups reflects the capability of SynActive to restrict the expression of the PSD95-FLAG bait to the subset of CA1 synapses subjected to potentiation after CxFC.

To gain insight into the potentiation-specific changes in the PSD-95 interactome, we compared samples from mice expressing SA-PSD95-FLAG exposed to the learning phase of CxFC with those from mice expressing Cnst-PSD95-FLAG kept in their home cage. We reasoned that this design would allow us to isolate differentially expressed proteins at potentiated synapses in a behavioral learning situation by comparison with constitutive components of all synapses in a behavioral baseline condition.

A few methodological considerations are worth underlining. The two constructs encoding PSD95- FLAG were delivered via AAV microinjection in the CA1 area, which restricted their expression to a subset of neurons in the dissected hippocampus. We chose not to use tandem affinity purification (TAP^14^) which, in principle, could improve the signal to noise ratio of IP, reasoning that the exquisite expression specificity of the proteomic bait for potentiated synapses over-compensates the signal to noise improvement that TAP would provide. In this regard, we adopted gentle protein extraction conditions to prevent disruption of the large complexes formed by PSD-95 and its indirect interactors. The validity of this approach is confirmed by the consistency between MS data and Western blotting validation for Arc and rpS6 proteins which are, indeed, indirect interactors of PSD-95.

On the other hand, including a group of animals that expressed no IP bait as a negative control enabled us to reduce the probability of artifactual findings generated by non-specific interactions with the anti-FLAG-coated IP beads. Remarkably, some of the 244 non-specific binders identified by this negative control were included in published datasets employing both similar (i.e. FLAG-based IP^14^) or radically different (i.e proximity labeling^41^) enrichment techniques, and 229 of them are contained in the comprehensive synaptome dataset published by Sorokina et al.^42^. While we cannot exclude that some of the non-specific binders identified by our FLAG-based method and contained in datasets from alternative approaches are true PSD-95 interactors, we opted for a high-stringency criterion and removed dubious entries from the analysis. In addition, we studied only proteins which showed significant enrichment in IP samples from SA or Cnst groups compared to Ctrl IPs. This was combined with dataset filtering based on available annotations on the cellular compartment to remove from the analysis proteins that are known not to be expressed in dendrites nor synapses.

Finally, we did not filter out proteins contained in the SA or Cnst datasets based on their significant enrichment in the IP vs. input comparison because this would have led to false negatives. Indeed, IP of FLAG-tagged PSD-95 selects the subset of units of a given protein that are interacting (either directly or indirectly) with the bait, while the input contains the entire cellular content of the same protein, including units that are not localized at synapses. If the latter population is larger than the former, no significant enrichment is observed, even if a biologically meaningful change in the interaction actually exists.

A direct comparison of filtered IP results from SA and Cnst groups was performed after normalizing each protein to the total abundance of the PSD-95 bait. We deemed this operation more correct than normalizing to the amount of PSD95-FLAG alone because the macromolecular complexes that assemble around this protein^54^ would contain both FLAG-tagged and untagged PSD-95, along with their interactors, that would be both pulled down by IP. For this reason, our datasets describe potentiation-induced changes in the stoichiometry of interaction between PSD-95 and its synaptic partners, rather than measuring differences in the total synaptic content of a given protein. This operation showed very large increases in the number of units of a given protein interacting with PSD- 95 after learning-induced synaptic potentiation, with a maximum log_2_(fold change) of 9.95 for the metabolic enzyme Ndufa5, and an average log_2_(fold change) of 4.37.

Our approach confirmed that induction of potentiation causes an average 4.37-fold increase in the synaptic abundance of AMPA receptor subunits^55,56^ and similarly affects kinases responsible for signal transduction and posttranslational modification of proteins such as PKA^57^, Rock2^58^, PI3 kinase^59^, CaMKIIα/β^60^. On the contrary, the average increase in the stoichiometry of interaction between PSD-95 and NMDA receptor subunits was 2.80-fold. This suggests that synaptic localization of NMDA receptors does not follow a 1:1 ratio with AMPA receptors, which can represent a possible molecular substrate for occlusion^61,62^ of subsequent rounds of potentiation. Of note, the fold change of Arc interaction in SA versus Cnst groups was also below the average value. This can be related to the proposed migration of this protein outside of potentiated synapses to mediate inverse tagging of non-potentiated synapses^63^ for endocytosis of AMPA receptors, a process that has been proposed to be involved in homeostatic synaptic scaling^64^.

Network analysis restricted to physically interacting proteins followed by categorization into main functional types showed that proteins enriched at potentiated synapses include receptors, ion channels, scaffolding proteins and kinases that are known to interact with PSD-95^14^. Most notably, the macromolecular complex hinged on PSD-95 also included a large cluster of interactors involved in protein translation including ribosomal proteins themselves. This agrees with evidence showing relocalization of polysomes^65^ and mRNAs^66^ at dendritic spines upon synaptic potentiation and with the well-known requirement of protein synthesis for maintenance of LTP. The SynActive-based approach allowed isolating individual components of the translational machinery enriched at synapses in response to behavioral learning and highlighted its molecular complexity. An equally interesting cluster contains components of vesicles, including several members of the SNARE family, which indicate increased trafficking required to add new structural components at potentiated synapses, reminiscent of the AMPA receptor exo- and endocytosis cycles involved in synaptic plasticity.

Finally, it is tempting to speculate that the increased abundance at potentiated synapses of mitochondrial proteins such as mt-Nd1 and mt-Nd2, as well as the electron transport chain metabolic enzyme Ndufa5 of as indirect interactors of PSD-95 is a sign of mitochondria remodeling and relocalization around dendritic spines^67^.

The proteins contained in the PSD-95 interactome were common to potentiated and total CA1 synapses. This indicates that increased synaptic transmission relies on quantitative, rather than qualitative, changes in the synaptic proteome, a principle which could be inferred from imaging and single-candidate studies^2,9,68^. The combination between state-dependent expression of a genetically encoded probe with mass spectrometry we adopted allowed its demonstration on a synaptome scale and is consistent with the qualitative similarity among the proteomes of the hippocampal trisynaptic circuit^15^.

Our results demonstrate that the SynActive approach can be successfully applied to quantitatively assess changes in the synapse proteome associated with potentiation triggered by in vivo behaviorally relevant stimuli. This is a significant advancement in comparison to previous studies, which qualitatively described the average PSD-95 interactome^13,14^, regardless of the synapse activation status.

A possible limitation of our study is the use of PSD-95 as bait. This protein is a central organizer of the postsynaptic structure surrounded by hundreds of interactors (^13,14^, and our data) that assemble into large macromolecular complexes held together by non-covalent bonds. As a consequence, direct but weakly bound interactors may be lost during the IP process, while some strong but spurious interactions may form after tissue lysis and loss of cellular compartmentalization. However, this is a common issue of all IP-based studies; in addition to a careful choice of the Co-IP conditions, as a compromise between stringency and gentleness, we minimized its impact on the validity of our data by adopting stringent dataset filtering criteria. A possible way to overcome this limitation is the use of proximity labeling-based tools^69,70^, which would covalently tag both stable and transient neighbors of the probe in situ. This will be part of our future efforts.

A second point to consider is the half-life of PSD-95 which, using Halo tagging, has been demonstrated to be between 5 and 7 days in the CA1 area^71^. Our protocol allows Cnst-PSD95-FLAG to reach steady-state expression, virtually resulting in its expression at all synapses of transduced neurons and mixing with endogenous PSD-95. On the other hand, SA-PSD95-FLAG has a much shorter time window for expression and localization at potentiated synapses where it will mix with the endogenous pool of PSD-95. PSD-95 forms large multimeric complexes^54^ which, in our case, may contain both endogenous and FLAGged units; however, the immunoprecipitation approach allows to select only PSD95-FLAG containing complexes, resulting in potentiated synapse-specific retrieval, regardless of the presence of endogenous PSD-95 synthesized prior to the induction of synaptic potentiation. This is also supported by the low baseline expression level of the construct in the absence of induction of synaptic potentiation by specific behavioral tasks (i.e., kept in their home cages).

From a conceptual point of view, the synaptic protein supercomplexes that support memory storage are constantly being turned over, translocated, substituted, and replaced^1,72^. Thus, future studies will go beyond a single time point to capture the evolution of the synaptic proteome at different moments during memory encoding and retrieval. The results presented here represent a proof of concept that highlights the potential of SynActive-based proteomics probes, as a strategy to investigate “memory synapses” in their dynamic time-dependent state^72,73^, and to compare the molecular correlates of synaptic potentiation in different phases of learning and memory (e.g., acquisition of conditioned behaviors and their recall), as well as in different types of behavioral tasks, in physiology or in different disease models (**Suppl. Fig. 15**).

For instance, SynActive could be used to control the expression of Halo-tagged proteins and combined with the DELTA approach to investigate protein turnover specifically at potentiated synapses^74^. Furthermore, the specificity for the potentiation status of SynActive can be combined with temporal control over bait expression (e.g., via conditional expression or doxycycline dependency), as well as with transgenic mouse lines expressing fluorescent synaptic reporters^75^, in future developments of our approach, in order to gain cell-type specificity and pinpoint the changes in the synaptic proteomes occurring at specific sub-phases of memory encoding and recall. The relevance of this aspect is, indeed, supported by the different distributions of potentiated synapses along the dendritic tree of CA1 pyramidal neurons during different phases of contextual fear conditioning learning and memory recall^18^.

We envisage that our method will aid the discovery of new molecules serving information storage at synapses in both physiological and pathological conditions, as well as in shifting the perspective of studies on the physical substrates of memory, i.e. engrams, from a cellular^39^ to a synaptic^27,76^ perspective.

## Supporting information

Supplementary file 1 - Cytoscape network

## ACKNOWLEDGMENTS

The research was funded by SNS institutional funds to AC and MM, MUR (Ministero Università e Ricerca) Progetti di Rilevante Interesse Nazionale (PRIN) 2017 project 2017HPTFFC to AC, PRIN 2022 project 2022MTR4M8 to AC and MM, Human Brain Project SGA2 (785907) to AC, NextGenerationEU – PNRR EBRAINS-Italy M4C2 (Project IR0000011) to AC, Next Generation EU – PNRR Tuscany Health Ecosystem (THE), Spoke 8, Project ECS_00000017 (MUR Directoral Decree n.1055 , 23/06/2022) to AC and MM, MUR Fondo Ordinario Enti (FOE D.M. n.789, 21/06/2023) – CNR-EBRI framework agreement to AC, and CNR project NutrAge (DSB.AD005.225) to MM.

We gratefully acknowledge Alessandro Viegi, Antonella Calvello, Vania Liverani for technical, infrastructural and organizational assistance; Francesca Biondi for assistance with the animal house; Rocco Pizzarelli (EBRI) for technical help in some initial experiments; Alessandro Cellerino (SNS) for productive and insightful discussions. We acknowldege support from the FLI Core Facility Proteomics. The FLI is a member of the Leibniz Association and is financially supported by the Federal Government of Germany and the State of Thuringia.

## AUTHOR CONTRIBUTIONS

AC, FG, and MM conceived the study and designed experiments. AO designed and supervised the proteomics experiments. IH performed mass spectrometry analyses. AC, MM, and SM secured grants supporting the project. AJ and FG prepared gene constructs and performed Western blotting and immunoprecipitations. LZ produced AAVs for in vivo gene transduction. AJ, MDC, MM, and SM performed in vitro validation experiments. MDC, AF, and MM performed in vivo AAV injections and validation experiments. FCL supported in vivo experiments. MM, IA, and AO performed proteomic data analyses. AC and MM wrote the manuscript.

---

**Supplementary Figure 1.**
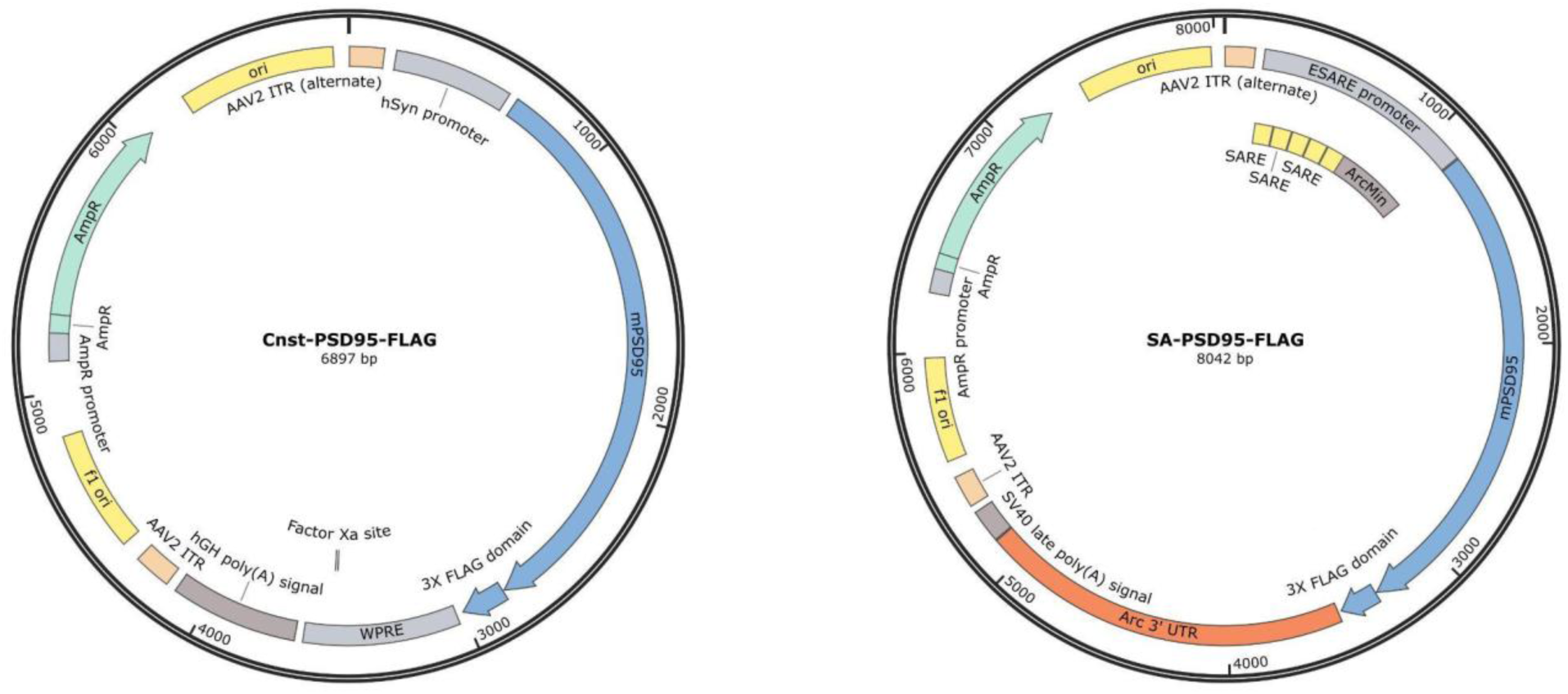
Maps of plasmid constructs used in this study.

**Supplementary Figure 2.**
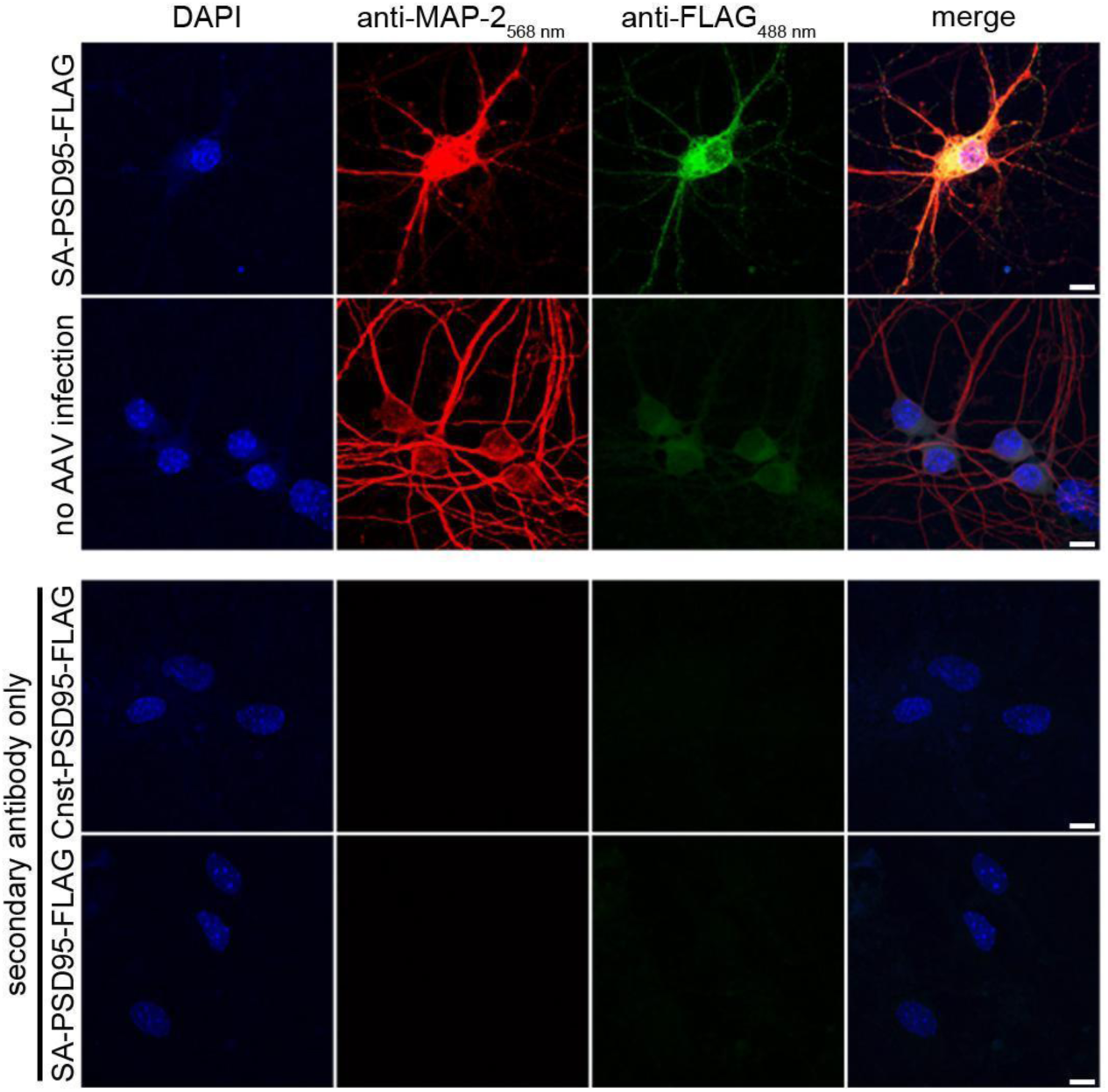
Technical controls for immunofluorescence on primary neuronal cultures. *Upper panel*, absence of FLAG immunoreactivity in primary cultures that did not receive infection with any AAV, while normal immunoreactivity for the endogenous protein MAP-2 was present. *Lower panel*, when primary antibodies (i.e. anti-FLAG and anti-MAP-2) were omitted, no immunoreactivity for FLAG and MAP-2 could be detected despite incubation with the corresponding anti-rabbit AlexaFluor-488 and anti-chicken AlexaFluor-568 secondary antibodies. Scale bars correspond to 10 μm.

**Supplementary Figure 3.**
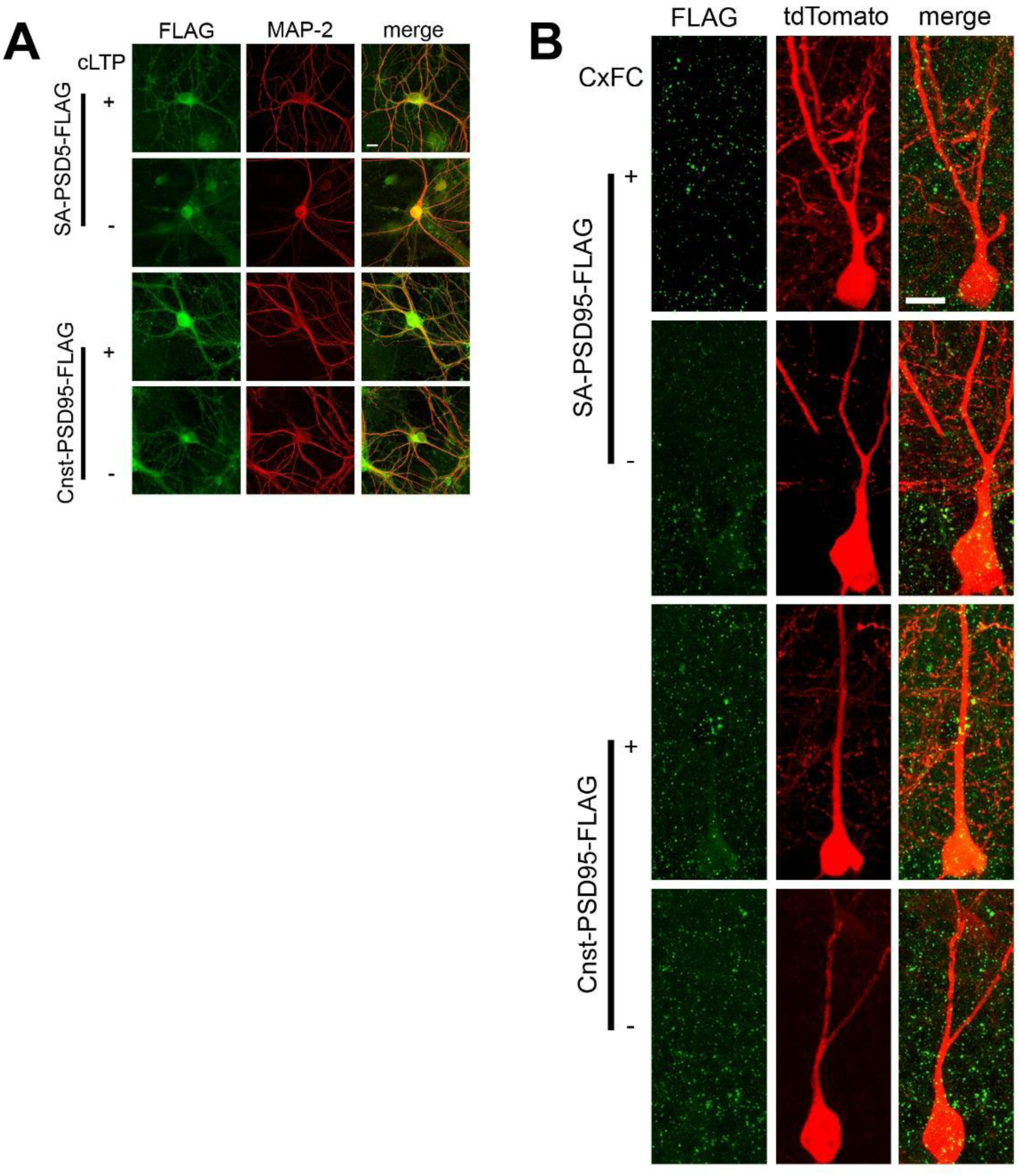
Representative images from **A)** primary hippocampal neuronal cultures and **B)** sections comprising the CA1 area showing that in vitro expression of SA-PSD95-FLAG (SA) and Cnst-PSD95-FLAG (Cnst) occurs in both dendrites and somata, while, in vivo, PSD95-FLAG displays a punctate pattern concentrated in dendrites. For panel B, tdTomato was sparsely transduced, thus leading to the visualization of the cell body and processes of a subset of all PSD95- FLAG-expressing neurons. This explains why FLAG-immunoreactive puncta can be observed also outside of tdTomato-positive neurons. Scale bars correspond to 20 μm.

**Supplementary Figure 4.**
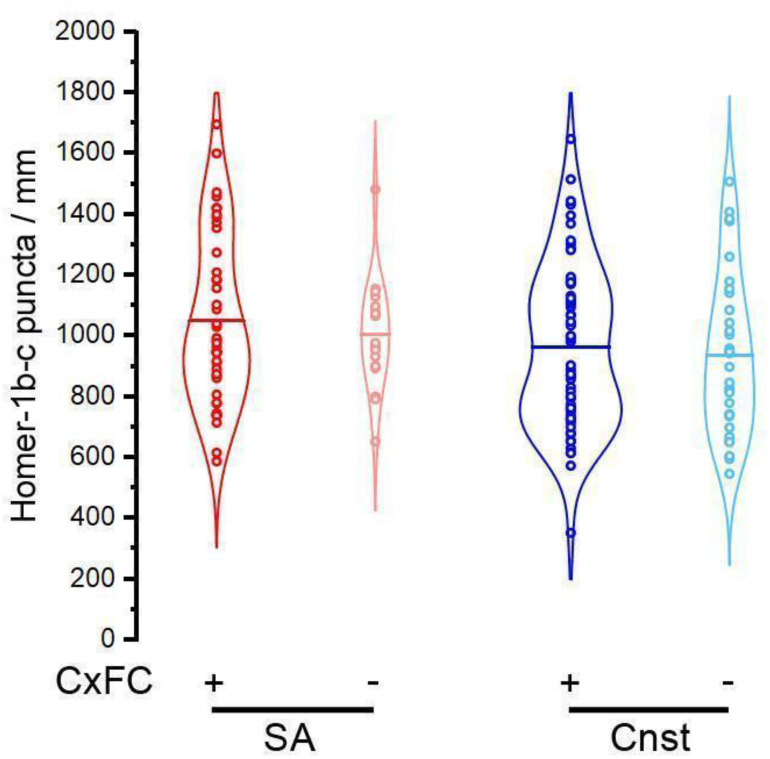
Unaltered Homer-1b-c positive synapse density after AAV-mediated in vivo expression of SA-PSD95-FLAG or Cnst-PSD95-FLAG. No significant difference in the density of Homer-1b-c immunoreactive synaptic puncta between experimental groups (ANOVA-2, SA vs Cnst, F_1,158_=2.795 p=0.097; CxFC+ vs CxFC-, F_1,158_=0.597 p=0.441; interaction, F_1,158_=0.0562 p=0.813; SA+CxFC, n=44; SA-CxFC, n=18; Cnst+CxFC, n=66; Cnst-CxFC, n=33 dendrites).

**Supplementary Figure 5.**
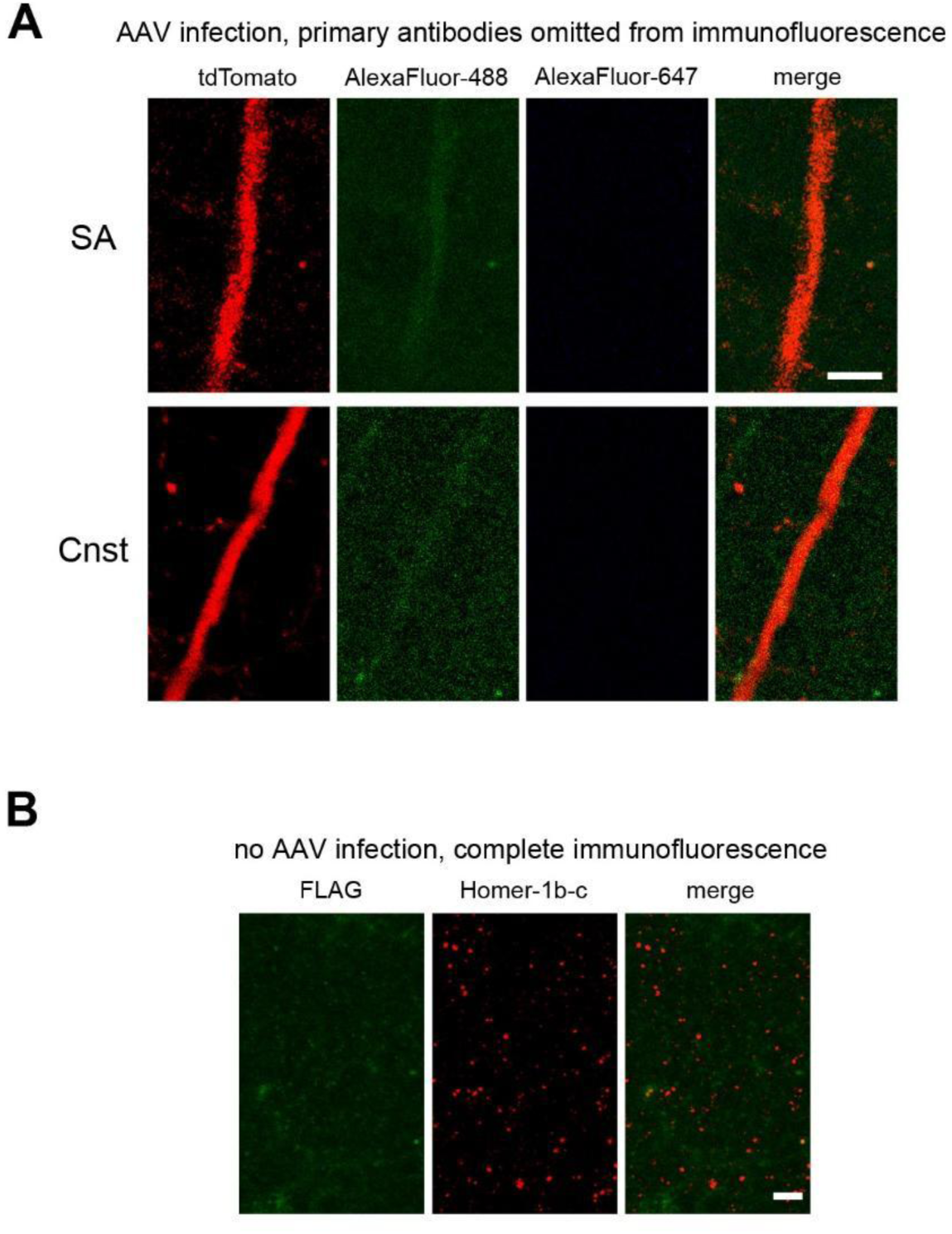
Technical controls for immunofluorescence on brain sections. **A)** No FLAG and Homer-1b-c signals could be detected in hippocampi from mice expressing SA- PSD95-FLAG and exposed to contextual fear conditioning, as well as from mice expressing Cnst- PSD95-FLAG, when the corresponding primary antibodies were omitted from the immunofluorescence reaction; the fluorescence associated with AAV-expressed tdTomato is present, as expected. **B)** Absence of FLAG immunoreactivity in hippocampi from mice that received no AAV infection and were processed using the same immunofluorescence procedure as in panel A; endogenous Homer-1b-c could be detected as expected. Scale bars correspond to 5 μm.

**Supplementary Figure 6.**
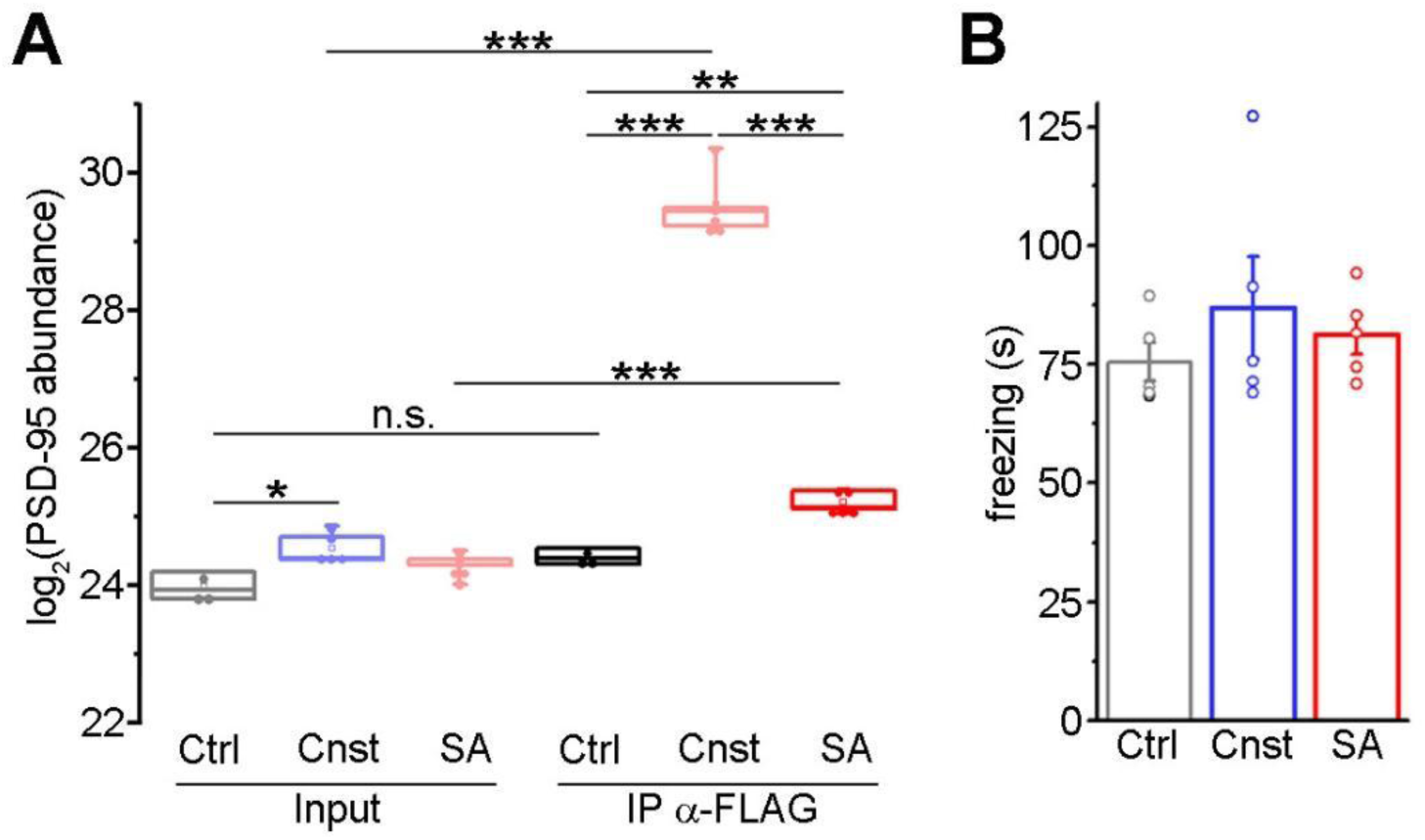
Impact of AAV-mediated expression of SA- or Cnst-PSD95-FLAG on PSD-95 expression and on contextual fear conditioning behavioral response. A) Constitutive expression of PSD95-FLAG using the synapsin promoter (Input-Cnst) led to a slight increase in the total levels of PSD-95 in comparison to mice that do not express PSD95-FLAG (Input-Ctrl), while no significant increase could be detected for mice expressing SA-PSD95-FLAG (Input-SA). FLAG immunoprecipitation (IP) resulted in a significant enrichment in PSD-95 in samples for SA and Ctrl groups with respect to the background levels in Ctrl mice; besides, IP-Cnst samples contained significantly more PSD-95 that IP-SA samples (ANOVA-2, F_2,20_=351.48, p(sample type × group)<0.001, followed by Holm-Sidak post hoc test, Input-Cnst vs Input-Ctrl, p=0.031; Cnst-Input vs Cnst-IP, p<0.001; SA-Input vs SA-IP, p<0.001; IP-Cnst vs IP-Ctrl, p<0.001; IP-Cnst vs IP-SA, p<0.001; IP-SA vs IP-Ctrl, p=0.002; Ctrl-Input and Ctrl-IP, n=3; Cnst-Input and Cnst-IP, n=5; SA- Input and SA-IP, n=5). B) No significant differences in the total duration of freezing during the learning phase of contextual fear conditioning among the Ctrl, Cnst and SA groups (ANOVA-1 F_2,12_=0.648, p=0.541; n=5 per group).

**Supplementary Figure 7.**
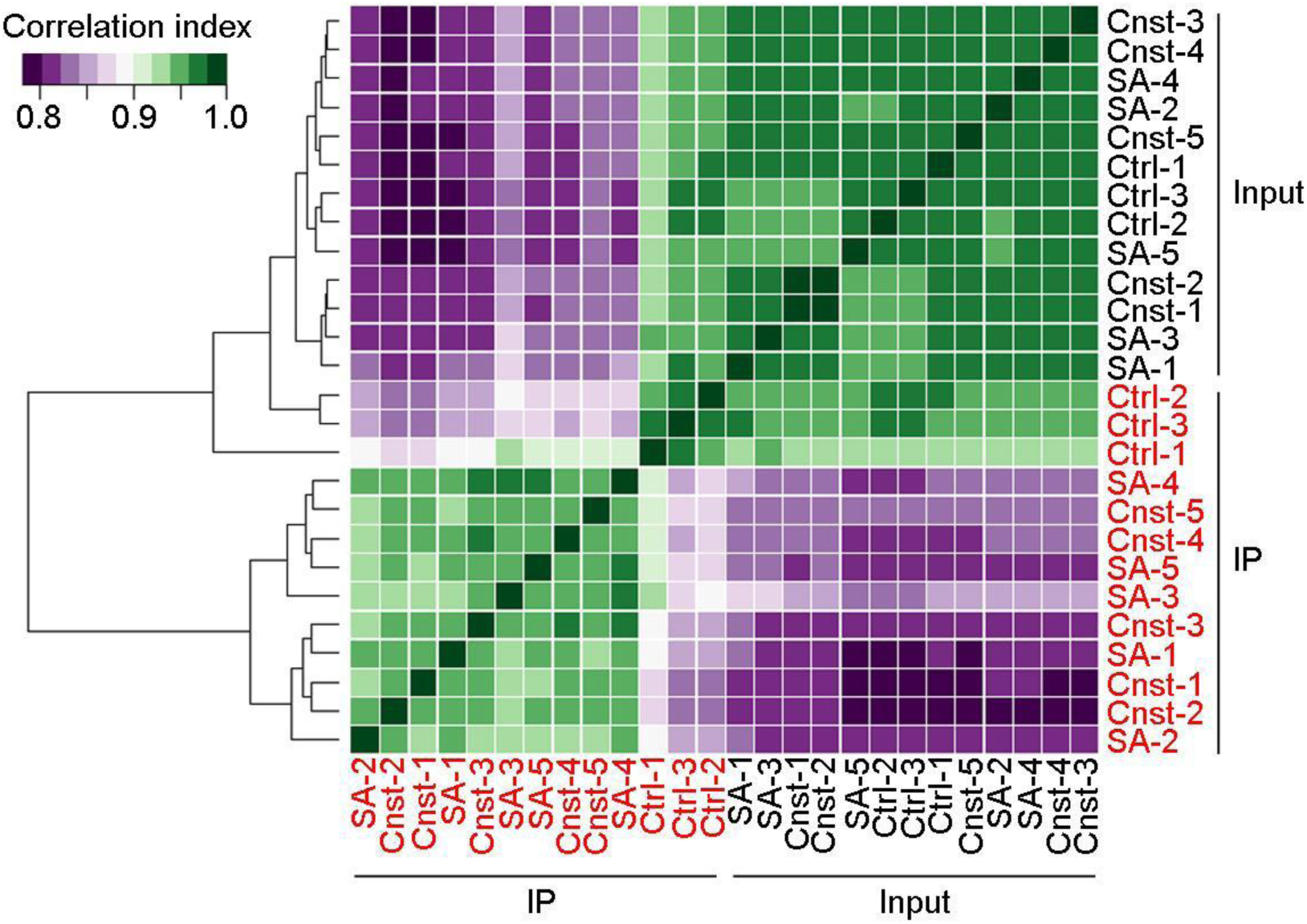
Analysis of global mass spectrometry data on samples from mice expressing SA- or Cnst-PSD95-FLAG. Correlation matrix and dendrogram for correlation indices among all samples, highlighting the separation between IP-MS and Input-MS data for Cnst and SA groups and the low IP-MS/Input-MS correlation for a given experimental group. On the contrary, Ctrl samples display a high IP-Input correlation.

**Supplementary Figure 8.**
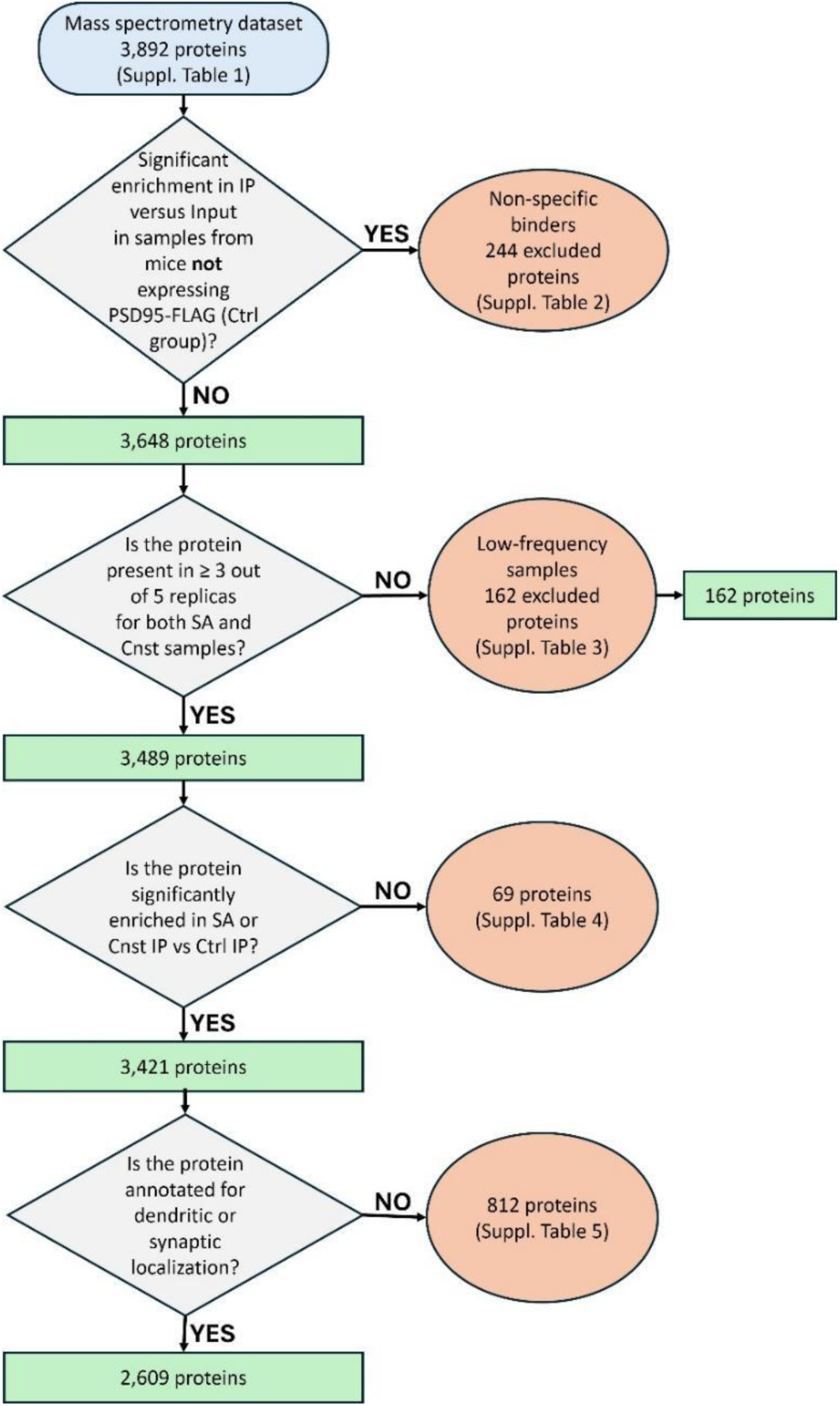
Flowchart describing the stepwise filtering approach of the raw MS protein dataset.

**Supplementary Figure 9.**
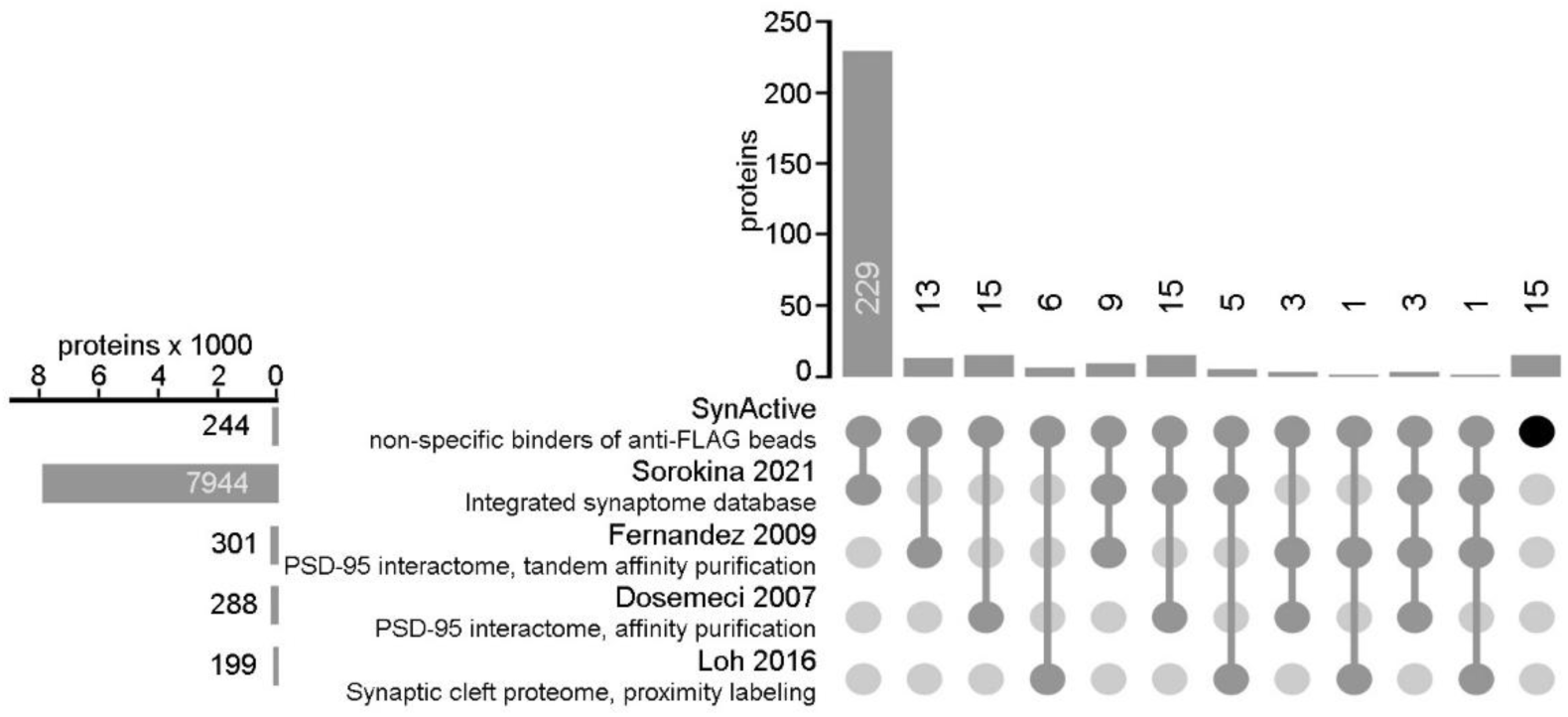
Comparison between the dataset of proteins non-specifically binding anti-FLAG-coated magnetic beads and published synaptomics datasets. This analysis highlights the presence of possible false positives in published datasets, with particular regard to those produced through anti-FLAG affinity purification, namely Fernandez 2009^14^ and Dosemeci 2007^13^. The intersection between our bona fide “non-specific binders” dataset and that by Loh 2016^41^, which was obtained via proximity labeling, indicates that even though some entries were classified as false positives in our anti-FLAG IP dataset, they could correspond to true synaptically localized proteins.

**Supplementary Figure 10.**
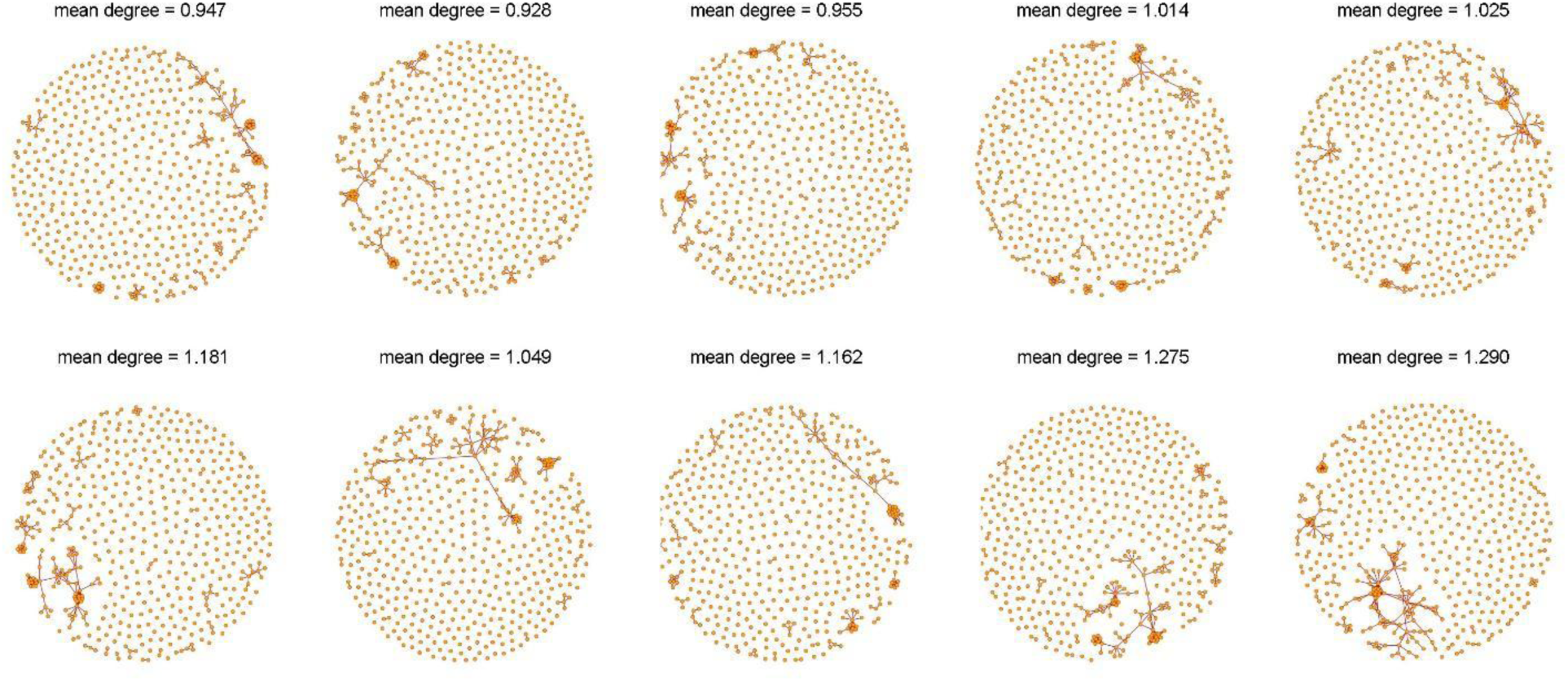
Examples of random networks. obtained from the same 513 nodes composing the interactome of PSD-95 enriched at potentiated synapses.

**Supplementary Figure 11.**
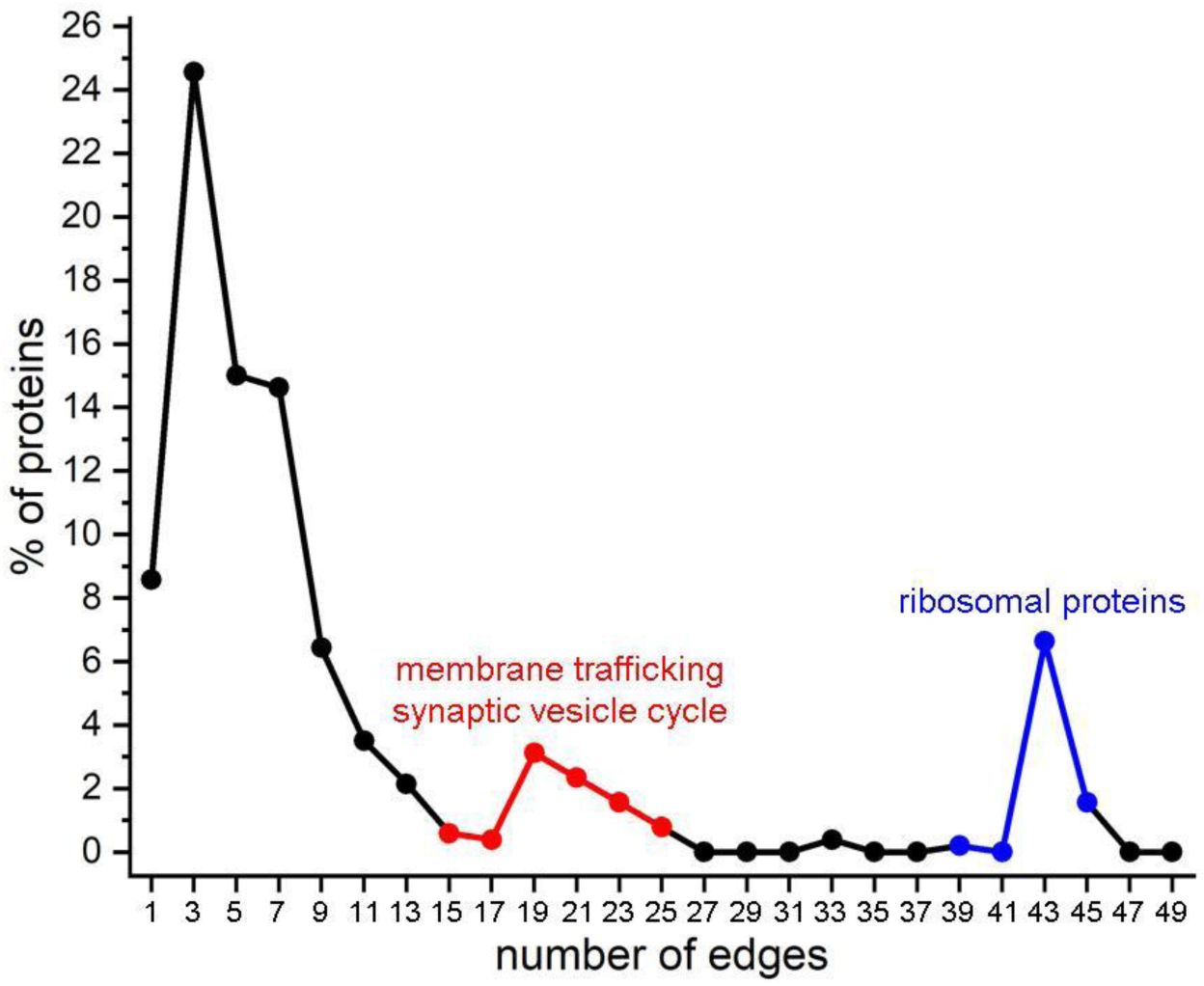
Percentage of proteins belonging to the PSD-95 interactome enriched at potentiated synapses showing a given connectivity degree.

**Supplementary Figure 12.**
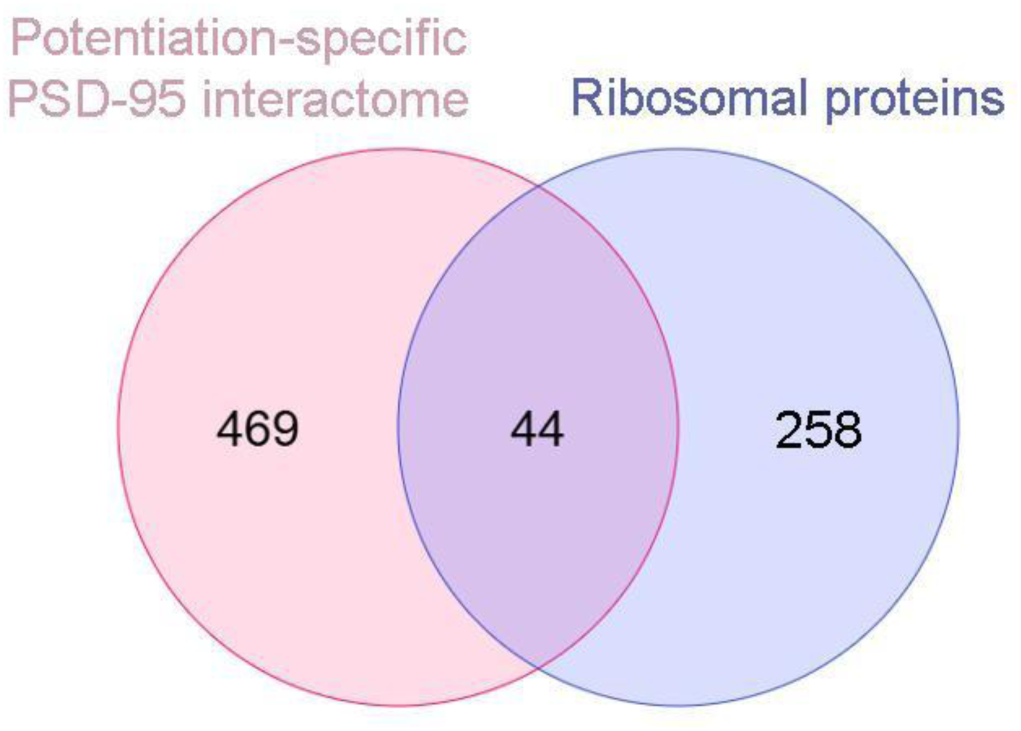
Venn diagram showing the relationship between ribosomal proteins enriched at potentiated synapses and those retrieved from UniProt by the query: "(28S OR 39S OR 40S OR 60S) AND "ribosomal protein" AND (organism_id:10090)".

**Supplementary Figure 13.**
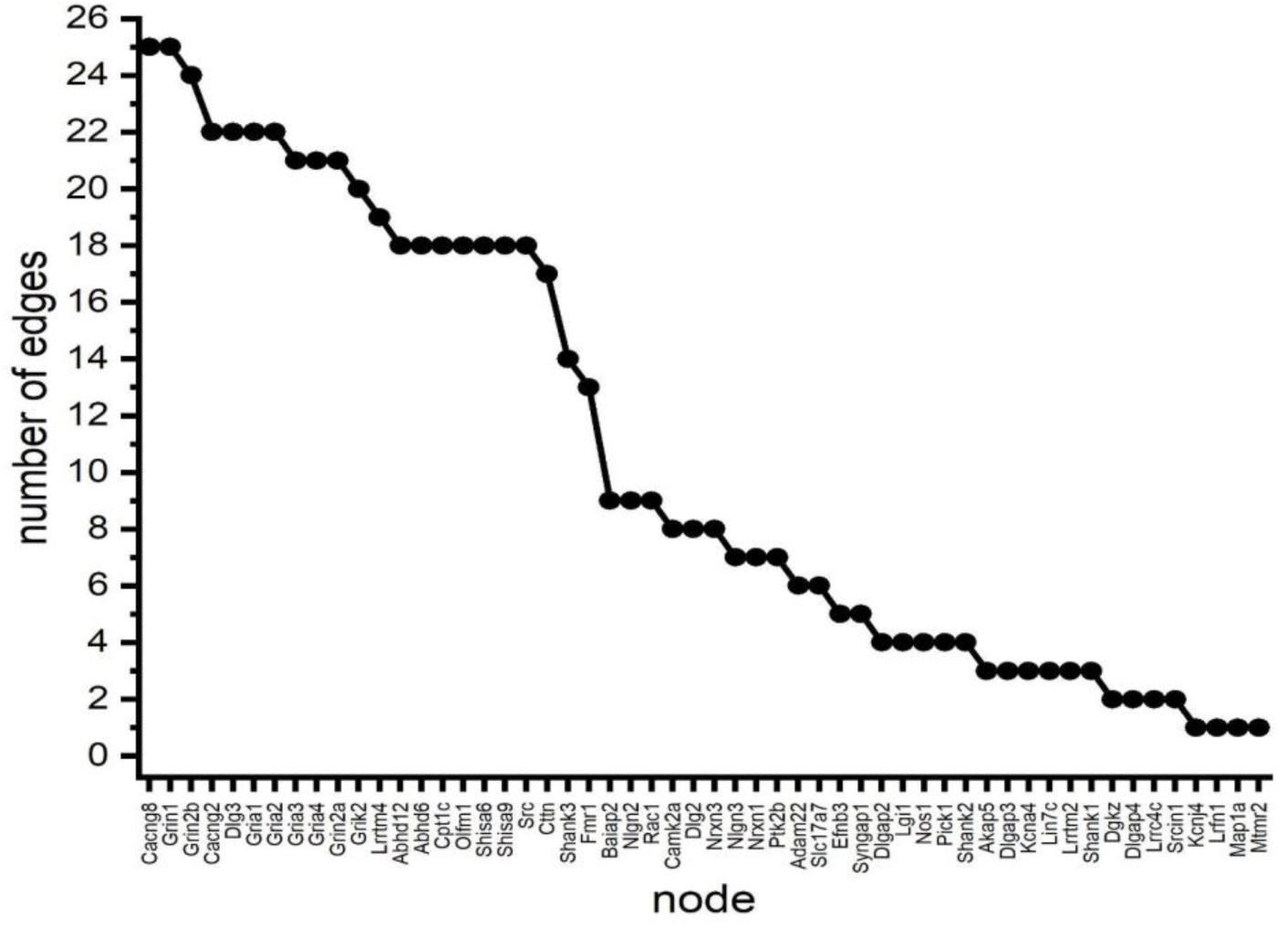
Number of connections for the 54 direct interactors of PSD-95 showing enrichment at potentiated synapses.

**Supplementary Figure 14.**
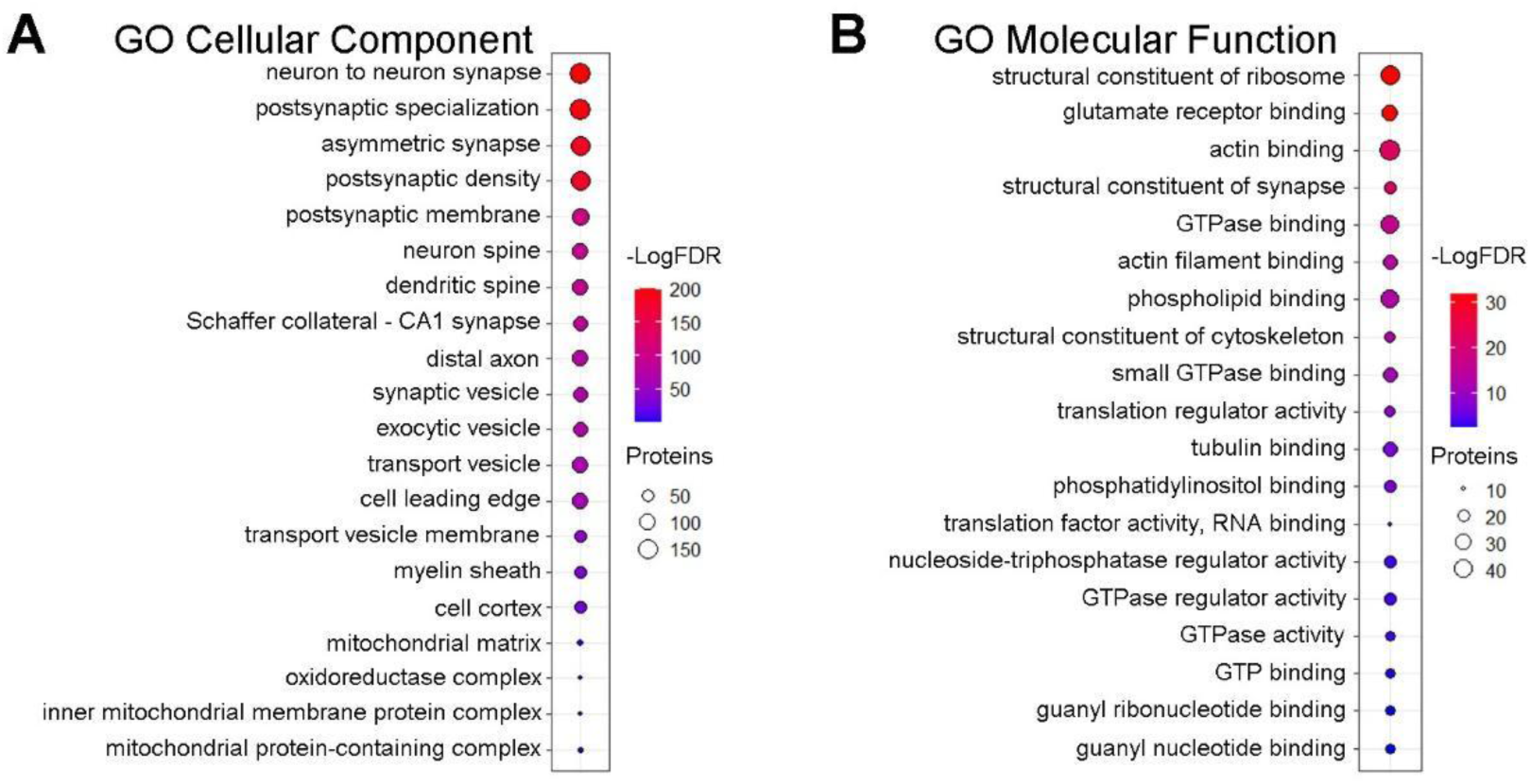
Gene ontology and pathways enriched in the PSD-95 interactome of potentiated synapses. **A, B)** Gene ontology (GO) Cellular Component and Molecular Function analyses for the nodes of the network shown in Fig. 6, ranked according to their odds ratio.

**Supplementary Figure 15.**
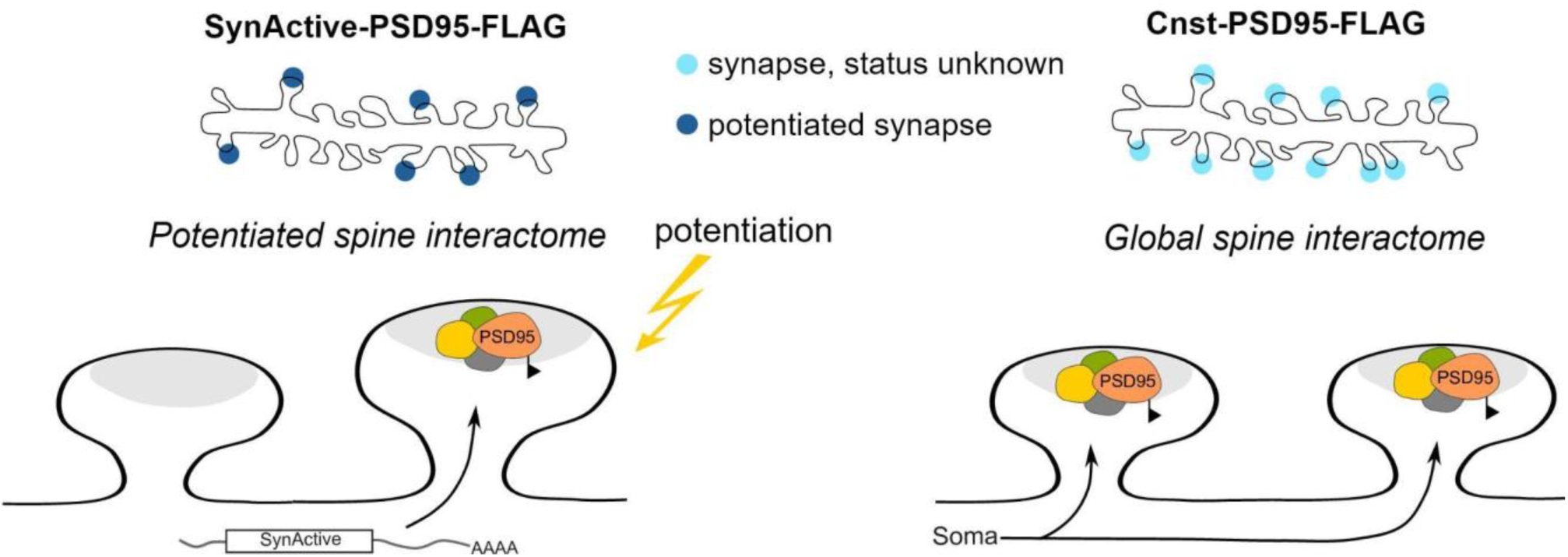
Schematic representation of the expression patterns of SynActive- PSD95-FLAG and Cnst-PSD95-FLAG constructs. The use of SynActive regulatory sequences enables the expression of PSD95-FLAG only at the subset of synapses undergoing potentiation via local translation. On the other hand, constitutive expression results in somatic translation of PSD95- FLAG, followed by its delivery to potentially all synapses of a neuron, regardless of their activation

